# Single cell lineage tracing reveals clonal dynamics of anti-EGFR therapy resistance in triple negative breast cancer

**DOI:** 10.1101/2023.04.04.535588

**Authors:** Simona Pellecchia, Melania Franchini, Gaetano Viscido, Riccardo Arnese, Gennaro Gambardella

**Author notes:** **These authors contributed equally to the work.** (Simona Pellecchia, Melania Franchini).

## Abstract

Epidermal growth factor receptor (EGFR)-targeted therapies have demonstrated variable and unpredictable clinical responses in triple negative breast cancer (TNBC). To elucidate the underlying mechanisms of this variability, we employ cellular barcoding and single-cell transcriptomics to reconstruct the subclonal dynamics of EGFR-amplified TNBC cells in response to afatinib, a tyrosine kinase inhibitor (TKI) that irreversibly inhibits EGFR. Integrated lineage tracing analysis revealed a rare pre-existing subpopulation of cells with distinct biological signature, including elevated expression levels of IGFBP2 (Insulin-Like Growth Factor Binding Protein 2). We show that IGFBP2 overexpression is sufficient to render TNBC cells tolerant to afatinib treatment by activating the compensatory IGF1-R signalling pathway. Finally, based on reconstructed mechanisms of resistance, we employ deep learning techniques to predict the afatinib sensitivity of TNBC cells. Our strategy proved effective in reconstructing the complex signalling network driving EGFR-targeted therapy resistance, offering new insights for the development of individualized treatment strategies in TNBC.

## Introduction

Triple-negative breast cancer (TNBC) is a highly heterogeneous and aggressive breast cancer subtype characterized by metastatic progression, poor prognosis, and the absence of targetable biomarkers (1–3). Neoadjuvant chemotherapy is initially highly effective on TNBCs but about 30%–50% of the patients rapidly develop resistance associated with higher mortality (1, 2, 4). Despite Epidermal Growth Factor Receptor (EGFR)-activating mutations and amplifications (≥ 5 copies) are uncommon in TNBC (3, 5–7), the majority of primary TNBCs exhibit enhanced expression of EGFR because of an increase in gene copy number (three to four copies) (7–12) (Supplementary Figure 01A-F), thus representing a valuable vulnerability for TNBC patients (13). However, unlike other tumour types where inhibition of wild-type EGFR by monoclonal antibodies or tyrosine kinase inhibitors (TKIs) is beneficial (14–17), EGFR-targeted therapies in TNBCs have shown variable and unpredictable clinical responses (1.7% to 38.7%) (18–22). Indeed, we and others (12, 20) found no significant correlation between EGFR status (*i.e.* copy number, mRNA, protein, or phospho-protein levels) and response to anti-EGFR therapies (Supplementary Figure 01G). Moreover, genomic variants that were found to be predictive of anti-EGFR resistance (23) in other tumour types are infrequent in TNBC patients (6, 24, 25) (Supplementary Figure 01H), suggesting that non-genomic mechanisms of resistance must be at play in TNBCs. As a result, the lack of predictive biomarkers of a response to anti-EGFR therapies in TNBC has hampered the translation of EGFR inhibitors in breast cancer.

Single-cell RNA sequencing (scRNA-seq) technologies have recently emerged as powerful tools to study intra-tumour heterogeneity (26, 27) and to reveal key genes that change in response to an external stimulus such as drug treatment (12, 28). However, when monitoring the emergence of drug-resistant cell populations, it is crucial to identify and compare how surviving lineages (*i.e.* clones) of cancer cells adapt to the treatment. Recently, bulk RNA sequencing and cellular barcoding, a technique that uses distinct DNA sequences to mark each cell, have provided the possibility of following cancer progression, metastasis dissemination, and cell differentiation at a single clone level (29). Hence, developing novel methods that couple DNA marking techniques with techniques measuring cellular states at single-cell resolution is essential to determine the different adaptive trajectories leading to drug resistance (28, 30, 31). Once these two techniques are coupled, *prospective* lineage tracing analyses can be used to study single clone dynamics during treatment to identify mechanisms driving drug adaptation, while *retrospective* analyses can be used to trace backwards in time and reconstruct pre-existing mechanisms of drug resistance by comparing the transcriptional states of tolerant and sensitive clones before the treatment (Figure 1A).

**Figure 1–.**
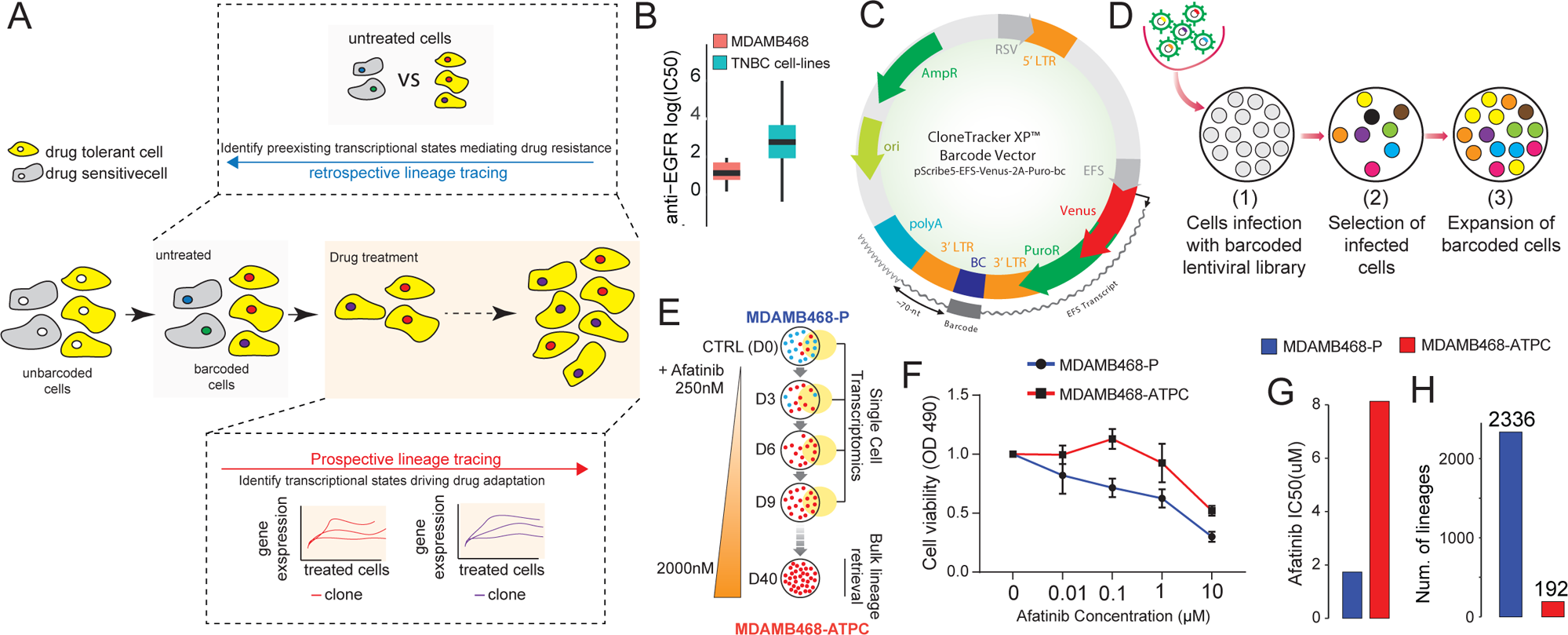
Platform for the identification of drug response biomarker genes with single cell lineage tracing. (**A**) An overview of strategies of lineage tracing. Once cells are marked (*i.e.* barcoded), they can be exposed to a selected drug concentration to drive the selection of resistant clones (yellow cells). Two types of analyses can then be performed. Prospective analyses where resistant clones (*i.e.* lineages) are followed over time during treatment to reconstruct driving mechanisms of drug adaptation. Retrospective analyses where resistant lineages are retrospectively mapped on the untreated condition to reconstruct pre-existing mechanisms of drug resistance. (**B**) Average log IC50 distribution of MDAMB468 cells and of all the other available TNBC cell lines to anticancer drugs targeting EGFR. (**C**) The Cellecta’s lentiviral construct contain a pool of random 48 base pairs (bp) barcodes located 70 bp far from ploy-A tail of the puromycin-resistance gene cloned downstream of a Venus fluorescent protein. (**D**) To enable lineage tracing in MDAMB468 cells, we seeded 50,000 cells and infected them with Cellecta’s lentiviral libraries at a multiplicity of infection (MOI) of 0.05. After five days of antibiotic selection, surviving cells were assessed by flow cytometry and subsequently expanded. (**E**) Scheme of the time series experiment performed on the barcoded MDAMB468 cell line. D0 are untreated parental MDAMB468 cells (MDAMB468-P) a few hours before afatinib addition, and D3 through D40 is the duration of the experiment where cells where subjected to incremental doses of afatinib ranging from 250 nM to 2000 nM. Cells at D40 were considered as afatinib tolerant persisted cells (ATPC). 10x Chromium sequencing was performed at Day 0 (D0), Day 3 (D3), Day 6 (D6) and Day 9 (D9) of 250 nM afatinib treatment. Quantseq-Flex to retrieve afatinib-tolerant lineages in bulk was performed at Day 40 with 2000 nM afatinib treatment. (**F**) Dose-response curve showing cell viability of MDAMB468-P and MDAMB468-ATPC cells following treatment with the indicated concentrations of afatinib. (**G**) Estimated IC50 for the dose response curve in (G). (**H**) Number of unique lineages present in the MDAMB468-P and MDAMB468-ATPC cell lines.

Here, we investigate the non-genomic mechanisms determining the cellular response to EGFR inhibitors in TNBCs. We integrate methods for cellular barcoding and single-cell transcriptomics to enable cell lineage tracing and explore the subclonal evolution of adaptation in an established preclinical model of TNBC (32, 33) in response to incremental concentrations of afatinib, a potent TKI that irreversibly inhibits both EGFR and HER2 receptors. Retrospective lineage tracing data analysis uncovered a pre-existing subpopulation of rare afatinib-tolerant cells displaying distinct biological features, such as elevated mRNA levels of IGFBP2 (Insulin-Like Growth Factor Binding Protein 2). We demonstrate by chemical and genetic manipulations that IGFBP2 overexpression is sufficient to render TNBC cells tolerant to afatinib treatment through activation of the compensatory IGF1-R signaling pathway. Prospective lineage tracing, on the other hand, highlighted additional adaptive mechanisms, including lysosome biogenesis, reactive oxygen species (ROS) homeostasis, and fatty acid metabolism. Finally, by linking reconstructed mechanisms of drug resistance with deep learning techniques we developed an algorithm to computationally predict afatinib response starting from the transcriptional status of TNBC cells. Our findings provide a new understanding of the intricate signaling network underlying EGFR-targeted therapy resistance in TNBC that can be helpful in devising novel strategies for TNBC patient stratification and therapeutic intervention.

## RESULTS

### 1. Single cell lineage tracing in EGFR-dependent TNBC cells

In this study, we use single-cell lineage tracing to identify novel predictive biomarkers of resistance to EGFR inhibitors in the human TNBC cell line MDAMB468. This cell line stands out among EGFR-wt TNBC cell lines due to its unique combination of EGFR gene amplification (>5 copies) and pronounced response to anti-EGFR therapies (Figure 1B) (32, 33), including afatinib (Supplementary Figure 02). As shown in Supplementary Figure 03, while other two TNBC cell lines may exhibit higher afatinib sensitivity, none of them harbour EGFR amplification, a key feature of EGFR-driven TNBC with potential for patient stratification. Furthermore, whole exome sequencing of MDAMB468 (34, 35) failed to identify mutations in genes linked to EGFR inhibitor resistance, such as KRAS and HRAS (23).

To enable single cell lineage tracing in these cells, we engineered them with Cellecta’s CloneTracker XP recorder technology. The Cellecta’s CloneTracker XP lentiviral barcode libraries (www.cellecta.com) are pooled expressed barcode libraries that enable the tracking and profiling of up to 10 million individual clones derived from a population of cells using either PCR or NGS techniques. As shown in Figure 1C, the lentiviral-based CloneTracker XP barcode libraries have two main functional elements: the reporter Venus protein and the drug resistance marker (PuroR), both expressed from a single promoter on a single transcript. The 48-nucleotide barcode cassette is embedded within the 3’-UTR sequence of the Venus mRNA and is located approximately 70 nucleotides upstream of the polyA. This design ensures that the barcodes are transcribed and can be captured during cDNA first-strand synthesis using standard oligo-dT primers employed in routine bulk and single-cell RNA-sequencing library preparation protocols. Indeed, after cell infection at low multiplicity of infection (Methods), antibiotic selection, and cellular expansion (Figure 1D), bulk RNA-sequencing analysis revealed that our founder population of MDAMB468 cells consisted of 2,336 unique barcodes (*i.e.* groups of related cells descended from a single clone).

Next, to induce the selection of afatinib-tolerant subpopulations, we subjected barcoded MDAMB468 cells to a gradient of increasing afatinib concentrations, ranging from 250 nM to 2000 nM (Methods). Over the course of the experiment, cells were collected at regular interval (*i.e.* every three days) for single cell transcriptomic profiling (Figure 1E). Following 40 days of selection, a stable subpopulation of cells emerged that could proliferate in a medium containing 2000 nM of afatinib. This resistant subpopulation, we named MDAMB468-ATPC (Afatinib Tolerant Persistant Cells), exhibited a 4-fold increase in afatinib IC50 values compared to the parental MDAMB468 population (Figure 1F,G). Barcode lineage retrieval from RNA bulk sequencing of MDAMB468-ATPC at 40 days revealed that only 192 out of the initial 2,336 clones (*i.e*., 8%) present in the parental MDAMB468 cell line survived to the afatinib selection process (Figure 1H).

### 2. Retrospective lineage tracing to elucidate pre-existing mechanisms of drug resistance of afatinib resistance in MDAMB468 cells

Next, we aimed to reconstruct non-genomic cellular states mediating the response to afatinib. To this end, we performed single-cell transcriptomic analysis on 4,101 cells harvested at four time points, day 0 (untreated), day 3, day 6 and day 9, following 250 nM of afatinib treatment (Figure 2A). By examining the transcriptional barcodes in individual cells, we were able to capture the transcriptional state of 448 distinct clones at day 0, while 186, 206 and 228 distinct clones were captured at day 3, day 6, and day 9, respectively.

**Figure 2–.**
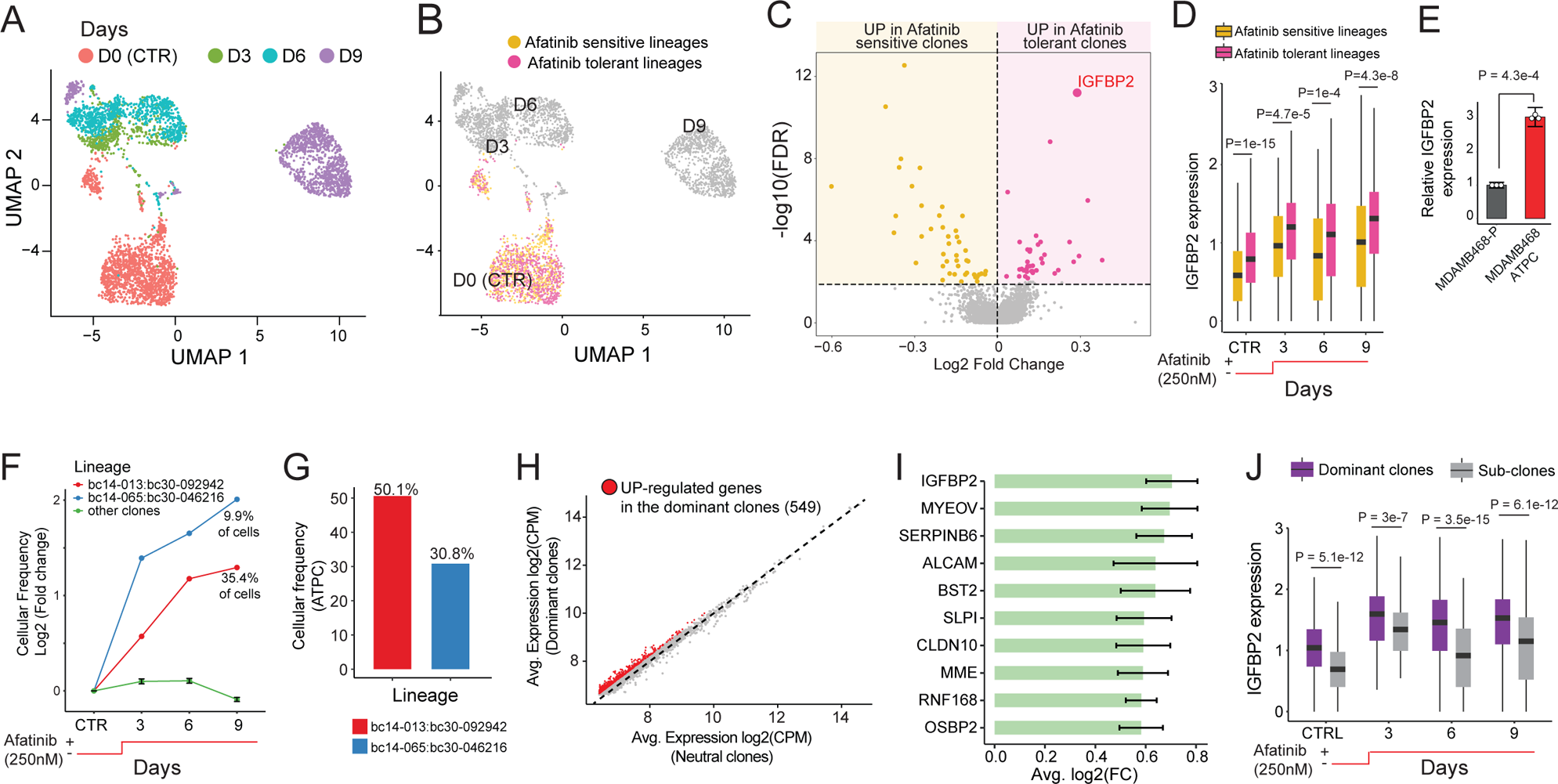
Retrospective and Prospective lineage tracing analysis. (**A**) UMAP representation of MDAMB468 cells colour-coded for the days of treatment. (**B**) Retrospective projection of the afatinib tolerant persisted lineages on a UMAP representation of untreated MDAMB468 cells at Day 0. (**C**) Differential expression between afatinib tolerant persisted lineages versus afatinib sensitive lineages on untreated MDAMB468 cells (*i.e.* cells from Day 0). (**D**) Expression distribution of IGFBP2 in afatinib tolerant persisted lineages and afatinib sensitive lineages over time. Statistical differences were estimated using a two-tailed Wilcoxon test. (**E**) IGFBP2 expression in MDAMB468-P and MDAMB468-ATPC cells measured by real-time quantitative PCR. The statistical difference was estimated using a two tailed t-test. (**F**) Cellular frequency of the afatinib tolerant clones bc14-013:bc30-092942 and bc14-013:bc30-092942 and all other clones at the sequenced time points (CTRL (Day 0), Day 3, Day 6, Day 9). (**G**) Cellular frequency of clones bc14-013:bc30-092942 and bc14-013:bc30-092942 in the MDAMB468-ATPC cell line. (**H**) Up-regulated genes identified by comparing cells belonging to the dominant clones to cells belonging to the neutral clones. (**I**) Average log2 fold change (FC) of the top ten up-regulated genes from (C). A 95% confident interval is reported. (**J**) Expression distribution of IGFBP2 in dominant and neutral clones over time. Statistical differences were estimated with a two-tailed Wilcoxon test.

We then retrospectively mapped the barcodes of 192 afatinib-tolerant clones present at day 40 onto the single-cell data of untreated MDAMB468 cells at day 0, as shown in Figure 2B. To assess whether afatinib-tolerant clones could be identified using conventional clustering approaches, we created a Uniform Manifold Approximation and Projection (UMAP) of D0 cells exclusively. As depicted in Supplementary Figure 04A, the afatinib-tolerant and -resistant clones are interspersed within the UMAP, implying a lack of distinct, specific expression patterns. To further corroborate this observation, we applied unsupervised clustering to the D0 cells (Supplementary Figure 04B). This analysis revealed the absence of any single cluster exclusively comprising tolerant or sensitive cells (Supplementary Figure 05C).

Next, we asked whether we could identify genes that were differentially expressed in untreated cells at day 0 by comparing cells belonging to afatinib-tolerant lineages with those belonging to the afatinib-sensitive lineages, i.e. those that were depleted after 40 days of continuous afatinib exposure. These genes, if present, should highlight pre-existing mechanisms of resistance to afatinib treatment. We found 374 genes that were differentially expressed between these two populations of cells (Supplementary Table 01). As shown in Figure 2C, among these genes, 221 were up-regulated in the afatinib-tolerant lineages (*i.e*, marker gene of resistance) and 153 were up-regulated in the afatinib-sensitive lineages (*i.e.*, marker genes of sensitivity). Gene Ontology Enrichment Analysis (GOEA) of the 221 up-regulated genes in the afatinib-tolerant cell population revealed several biological processes associated with resistance to EGFR inhibitors in other cancer types, such as oxidative phosphorylation and fatty acid metabolism (28, 36–38) (Supplementary Table 02). In contrast, the 153 genes up-regulated in the afatinib-sensitive cell population included EGFR and several genes related to the estrogen signalling pathway (Supplementary Table 02).

As shown in Figure 2C, among the 221 up-regulated genes in the afatinib-tolerant cell population, the most significantly upregulated gene was the Insulin-Like Growth Factor Binding Protein 2 (IGFBP2), which is a member of a family of six proteins specifically binding insulin-like growth receptors I and II (IGF1-R and IGF2-R). IGFBP2 has been recently implicated in the progression and metastasis of several tumour types (39, 40). Interestingly, as shown in (Figure 2D), the overexpression of IGFBP2 in afatinib-tolerant lineages is maintained during treatment, suggesting that this gene, among the others, could be required both in drug resistance initiation and maintenance. Indeed, higher expression of the IGFBP2 gene in afatinib-tolerant clones was further confirmed by qRT-PCR of MDAMB468-ATPC (Figure 2E).

Previous studies have shown that resistance mechanisms to anti-EGFR therapies involve the compensatory activation of signalling pathways that share downstream effectors with the EGFR signalling cascade, including IGF1-R, MET and the PI3K-AKT-mTOR signalling pathways (23, 41–44). This understanding, coupled with the knowledge that IGFBP2 is part of the IGF1-R pathway, prompted us to co-treat parental MDA-MB-468 cells with different concentrations of Afatinib and GSK-1904529A, a selective inhibitor of IGF1-R and Insulin Receptor (INSR). To gain a broader understanding of this dose-response relationship, we performed two independent experiments in triplicate, one in which afatinib varied from 1 to 16 μM (Supplementary Figure 05A,B) and another in which it varied between 31 nM and 1 μM (Supplementary Figure 05C,D). As shown in Supplementary Figure 05, the synergism between afatinib and GSK4529 is evident over a broad range of afatinib concentrations but is potent around 1 μM. These results support the existence of a pre-existing group of cells with enhanced activity of the compensatory IGF1-R signalling pathway could be sustained by IGFBP2.

In conclusion, these findings collectively emphasize the significance of our retrospective lineage strategy in identifying subtle distinctions that could otherwise be overlooked by conventional clustering methods based solely on gene expression analysis. Our strategy reviled IGFBP2 as a potential player in initiating afatinib resistance in MDAMB468 cells.

### 3. Prospective lineage tracing to reconstruct clonal expansion patterns in afatinib-tolerant MDAMB468 cells

Our previous analyses revealed that only 192 out of 2,336 clones (*i.e.* lineages) originally present in the parental MDAMB468 cell population became fully tolerant to afatinib treatment. To understand how these clones evolved over time, we computed at each sequenced time point the percentage of cells associated with each afatinib-tolerant lineage and plotted them over time, as shown in Figure 2F (and Supplementary Table 03). This analysis revealed that 190 of the 192 lineages were “neutral”, maintaining a constant relative population size (*i.e.* the same percentage) over time. Two clones, however, showed significantly increased relative size frequency, accounting for almost half (45%) of the total population after nine days of treatment (Figure 2F). Further analysis confirmed that these two clones became the “dominant” ones, comprising 81% (Figure 2G) of the drug-tolerant MDAMB468-ATPC cell line (*i.e.*, cells after 40 days of afatinib exposure).

Next, we sought to determine if the two dominant clones displayed distinct characteristics compared to the remaining 190 neutral clones, allowing their identification using conventional clustering approaches. To address this question, we generated UMAP visualizations of tolerant cells at each time point and clustered them based on their transcriptional similarities (Supplementary Figure 06). As seen in Supplementary Figure 06, no single cluster exclusively houses cells derived from either the dominant or neutral clones, except for cluster 5 of Day 0 tolerant cells, which however comprises only 12 cells.

To identify key genes driving the cellular expansion of the two dominant clones during the first nine days of afatinib treatment, we divided cells into two groups according to the behaviour of their clone of origin, *i.e.* “dominant” or “neutral”. Then, we ordered the two groups of cells along a linear pseudotime (pt) trajectory (45) from day zero to day nine and used time-course differential expression analysis (46) to identify genes that were up-regulated in the cells of the dominant clones (see Methods). This clone resolution analysis identified a total of 549 driver genes that were significantly up-regulated (FDR < 5%) in the dominant clones (Figure 2H and Supplementary Table 04), with IGFBP2 again being the most up-regulated gene (Figure 2I,J). These findings suggest that IGFBP2 could be a player in both resistance initiation and cellular expansion during afatinib treatment.

To test this hypothesis, we generated two novel cell lines from the parental MDAMB468 using CRISPRa and CRISPRi techniques: the MDAMB468-VPH-IGFBP2 (stably overexpressed IGFBP2) and the MDAMB468-KRAB-IGFBP2 (stably downregulated IGFBP2) (Supplementary Figure 07A,B and Methods). As shown in Figure 3A,B IGFBP2 knockdown significantly increased afatinib cytotoxicity while IGFBP2 overexpression significantly decreased it. Furthermore, IGFBP2 overexpression allowed cell colony growth even with a concentration of afatinib ranging from 2 to 10 μM (Figure 3C). These results were further validated in 3D cell culture, where IGFBP2 overexpression increased MDAMB468 spheroid growth at 2 μM afatinib exposure (Figure 3D). In contrast, IGFBP2 knockdown inhibited colony formation after 5 days of exposure to 1 μM afatinib (Figure 3E) and significantly inhibited MDAMB468 spheroid growth during afatinib treatment (Figure 3F). To investigate whether IGFBP2 could confer resistance to afatinib in other TNBC cell lines, we selected additional highly afatinib-responsive (Supplementary Figure 03) TNBC cell lines (HCC1806 and HDQP1) and again employed the CRISPRa system to generate cells stably overexpressing IGFBP2 (Supplementary Figure 07C). As shown in Supplementary Figure 08A,B IGFBP2 overexpression decreased afatinib cytotoxicity in both cell lines.

**Figure 3–.**
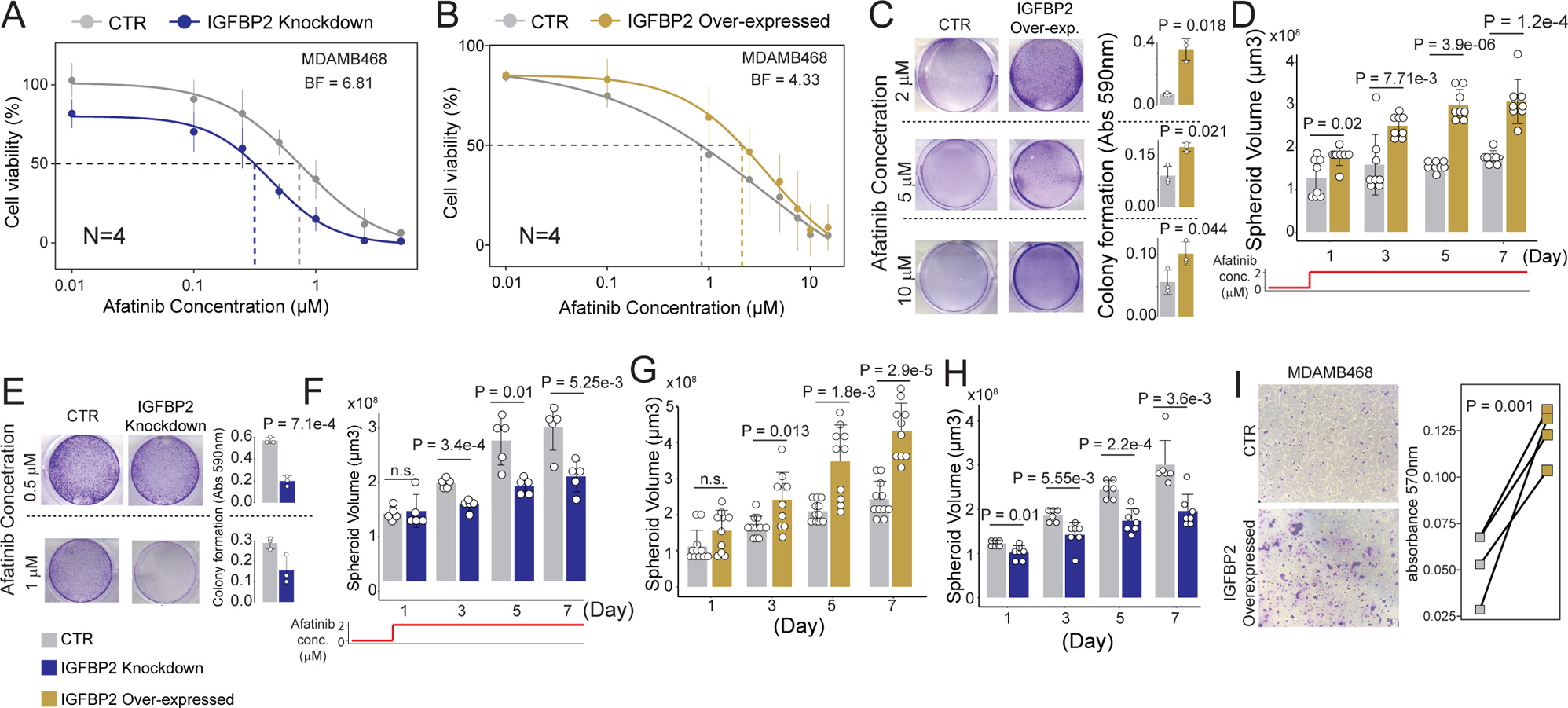
Identification of genes driving cellular expansion during afatinib adaptation. (**A**) Dose-response curve in terms of cell viability following treatment of MDAMB468 control (CTR) and IGFBP2-knockdown cells with afatinib at the indicated concentrations. Significance was assessed using Gaussian processes and BF is the estimated Bayesian Factor expressed in bits representing the difference between the two dose-response curves (Methods). A value of 6.81 corresponds to a very strong significant difference between the two dose-response curves (77). (**B**) Dose-response curve in terms of cell viability of MDAMB468 control (CTR) and IGFBP2-overexpressing cells following treatment with afatinib at the indicated concentrations. Significance was assessed as in (A). A value of 4.3 corresponds to a strong significant difference between the two dose-response curves (77). (**C**) Colony assay (representative images) for MDAMB468 control (CTR) and IGFBP2-overexpressing cells after 10 days of afatinib exposure at 2, 5 and 10 μM (left). Quantification is reported on the right. Experiments were performed in triplicate. (**D**) Spheroid volume growth of MDAMB468 control and IGFBP2-overexpressing cells with 2 μM afatinib (see Methods) over time. (**E**) Colony assay (representative images) for MDAMB468 control and IGFBP2-knockdown cells after 3 days of afatinib exposure at 0.5 and 1μM (left). Quantification is reported on the right. Experiments were performed in triplicate. (**F**) Spheroid volume growth of MDAMB468 control and IGFBP2-knockdown cells with 2 μM of afatinib over time. (**G**) Spheroid volume growth over time of MDAMB468 IGFBP2-overexpressing cells. (**H**) Spheroid volume growth over time of MDAMB468 IGFBP2-knockdown cells. (**I**) Transwell migration assay of MDAMB468 IGFBP2-overexpressing cells. Reported p values from panels C to I were estimated using two-tailed t-test.

Finally, to dissect the biological process sustained by IGFBP2 expression, we performed bulk 3’ RNA-sequencing of MDAMB468-VHP-IGFBP2 cells. Our analysis identified 856 transcripts that were significantly modulated by IGFBP2 overexpression (FDR < 0.05), with 373 were up-regulated and 483 down-regulated (Supplementary Table 05). GOEA of the 856 differentially expressed genes showed a strong enrichment for genes related to the PI3K-AKT axis/AMPK signalling (40, 47) but also many genes related to cell adhesion, growth, and migration (Supplementary Table 06). Indeed, IGFBP2 knockdown impaired spheroid growth of untreated MDAMB468 cells, conversely to its overexpression (Figure 3G,H). IGFBP2 overexpression also increases cell migration in MDAMB468 (Figure 3I) and additional TN cell lines such as HCC1806 and HDQP1 (Supplementary Figure 08C,D). Moreover, 88 of the 856 genes modulated by IGFBP2 overexpression were also found among the 549 driver genes upregulated in the dominant lineages. These 88 genes were primarily associated to cell growth, adhesion, and metabolic processes (Supplementary Table 07).

In conclusion, we have demonstrated the capacity of our cell lineage tracing strategy to enable the reconstruction of cell expansion at single-clone resolution, providing insights into the dynamic nature of drug resistance mechanisms. Our findings also reinforce the role of IGFBP2 as one of the players influencing afatinib adaptation in MDAMB468 cells.

### 4. Temporal analysis of afatinib-mediated transcriptional programs in MDAMB468 cells

To investigate the coordination of gene activity and transcriptional programs among drug-resistant sub-populations of cancer cells, we clustered the expression patterns of the 549 driver genes upregulated in drug-tolerant lineages along their reconstructed pseudotime trajectory (Methods). After clustering, four distinct patterns of transcriptional adaptation to afatinib treatment emerged (Figure 4A). Cluster one included genes with an “early” transcriptional response, increasing significantly after three days of treatment, while cluster two included genes with a “delayed” transcriptional response, increasing significantly after six days of treatment. Cluster three included genes that progressively downregulate in expression during afatinib treatment, while genes in cluster four exhibited a transient decrease in expression followed by a return to baseline levels.

**Figure 4–.**
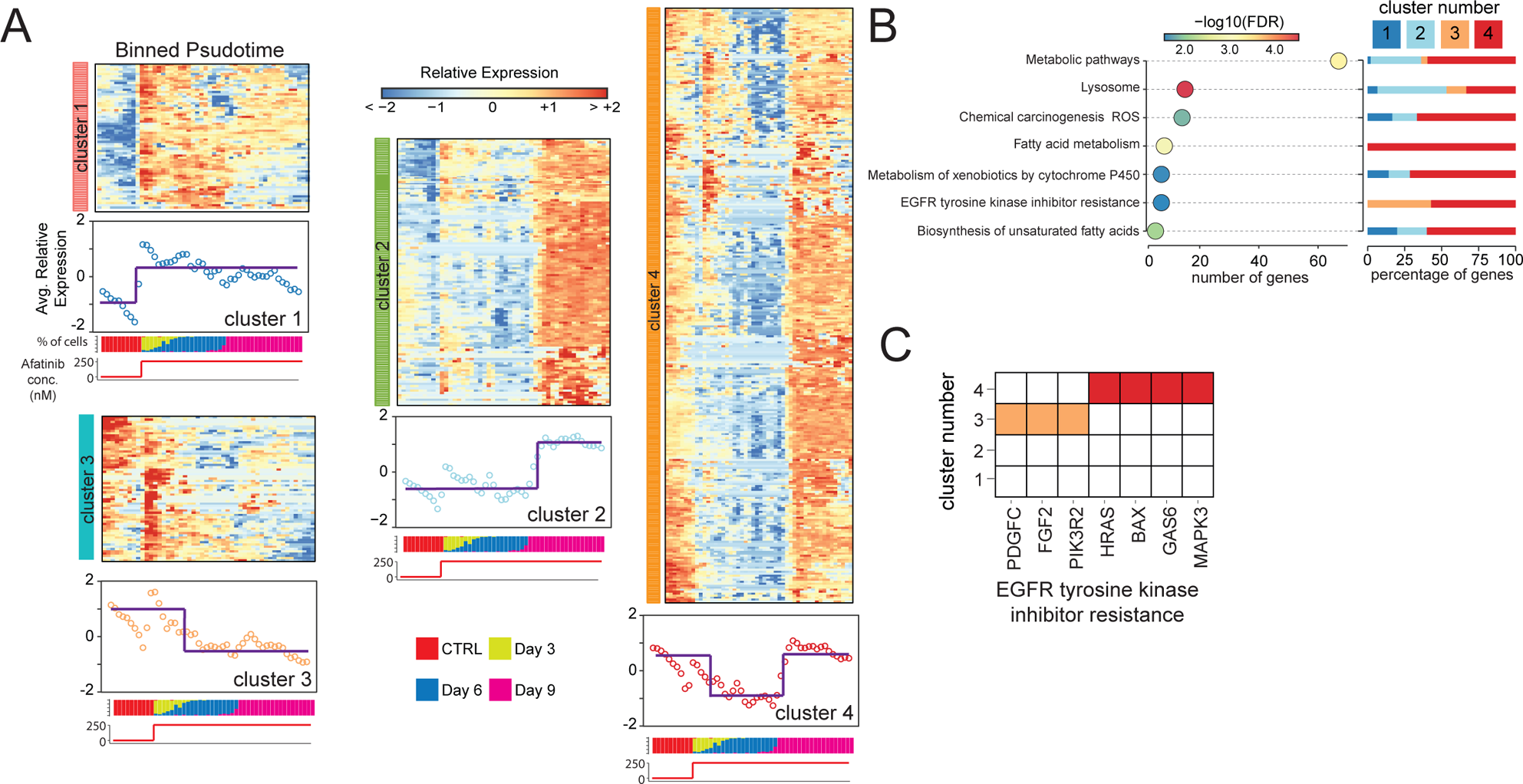
Gene expression dynamics in response to afatinib adaptation. (**A**) 549 driver genes clustered according to their expression. Cells are binned according to their reconstructed pseudotime. The average expression profile of the genes in the cluster and the cell composition of each bin as a percentage of cells in each timepoint is reported below each cluster. (**B**) Gene ontology enrichment analysis of 549 genes (bottom) and for each enriched term the percentage of genes belonging to a specific cluster of (A). (**C**) Genes in the enriched term EGFR tyrosine kinase inhibitor resistance and the cluster of (A) in which they fall.

Next, we employed time-resolved Gene Ontology Enrichment Analyses (GOEA) to decipher the sequential activation of the transcriptional programs driving afatinib adaptation. To this end, we first performed GOEA considering all 549 driver genes (Supplementary Table 08) and then reconstructed the specific activation time of each significant GO term by calculating the proportion of genes it included from each cluster. GOEA analysis of the 549 driver genes (Figure 4B) revealed several biological processes previously linked to drug resistance in other cancer types, including lysosome biogenesis, Reactive Oxygen Species (ROS) homeostasis, and fatty acid metabolism (48–54), as well as specific genes linked to EGFR resistance, such as PDGF-C (55), FGF2 (56, 57), PIK3R2 (58), HRAS (59), MAPK3 (60, 61), GAS6 (62), and BAX (63) (Figure 4C).

Interestingly, PDGF-C and FGF2 are PDGFR-α and FGFR ligands, respectively. They are well-established compensatory pathways activated to overcome EGFR inhibition (64, 65) often associated with the acquisition of mesenchymal features (64, 65) sustained by the AXL pathway (66, 67) whose activator is GAS6 (Figure 4C). Lysosome biogenesis and lipogenesis have been extensively studied for their involvement in drug adaptation to many anticancer drugs in multiple cancer types, including both standard chemotherapeutics and targeted therapy (48–54). Enhanced lysosomal biogenesis enables the sequestration of hydrophobic weak base compounds, thereby reducing the cytotoxic potential of a drug by limiting its availability at the site of action (48, 49). On the other hand, enhanced lipogenesis has been reported to contribute to drug adaptation by reducing drug uptake and participating in antioxidant cell defence by regulating membrane fluidity (50–54).

As shown in Figure 4B, a temporal program emerges, although most pathways exhibit genes distributed across all four clusters. For instance, lysosomal genes are particularly enriched in clusters one and two, suggesting a possible cellular early response to drug-induced damage. Conversely, genes associated with fatty acid metabolism, ROS, and EGFR TKI resistance (Figure 4B,C) exhibit expression profiles falling within clusters three and four, where gene expression is elevated from day 0 (i.e., untreated cells), suggesting that these pathways could be inherent characteristics of drug-tolerant cells that are either required in later stages of drug adaptation (cluster four) or not (cluster three).

In conclusion, these results support the hypothesis that during the afatinib response, along with the induction of xenobiotic detoxification mechanisms to reduce drug availability, tolerant clones activate compensatory pathways beyond IGF1-R to maintain pro-survival signals. This enables them to reduce Reactive Oxygen Species (ROS) production (68) while mitigating their damaging effects by producing antioxidants and facilitating Epithelial to Mesenchymal Transition (EMT) (65, 68).

### 5. Development of a deep learning approach to predict afatinib Sensitivity in TNBC

Our retrospective lineage tracing analysis on barcoded MDAMB468 cells identified 374 genes whose expression levels significantly influence afatinib responsiveness. This led us to question whether these genes could serve as effective predictive biomarkers for afatinib therapy in other TNBC cells. To address this question, we developed scASTRAL (<u>s</u>ingle-<u>c</u>ell <u>A</u>fatinib re<u>S</u>ponse of <u>TR</u>iple neg<u>A</u>tive ce<u>L</u>ls), a deep doublet learning approach (Methods) that utilizes both contrastive learning and Supported Vector Machine (SVM) to predict afatinib sensitivity from single cell data, leveraging the expression levels of the 374 marker genes identified in Figure 2C.

Contrastive learning is a machine learning approach that aims to construct an optimized embedding space where similar sample pairs are pushed closer and dissimilar ones farther apart. As shown in Figure 5A, scASTRAL uses as input cells whose transcriptional profiles are represented by the expression levels of the 374 marker genes of the afatinib response identified previously. The algorithm involves two main steps. In the first step, scASTRAL builds an embedding space where the cosine distance between cells belonging to clones of the same afatinib-response class (e.g., afatinib-sensitive or afatinib-tolerant) is minimized, while at the same time, the distance between cells belonging to clones of different afatinib-response classes is maximized. As illustrated in Figure 5B, this effectively separates cells based on their afatinib response. In the second step scASTRAL trains an SVM classifier on the embedded data to distinguish afatinib-sensitive and afatinib-tolerant cells with high accuracy. This classifier can then be used to predict the afatinib sensitivity of new single-cell data from other treatment-naïve TNBCs.

**Figure 5–.**
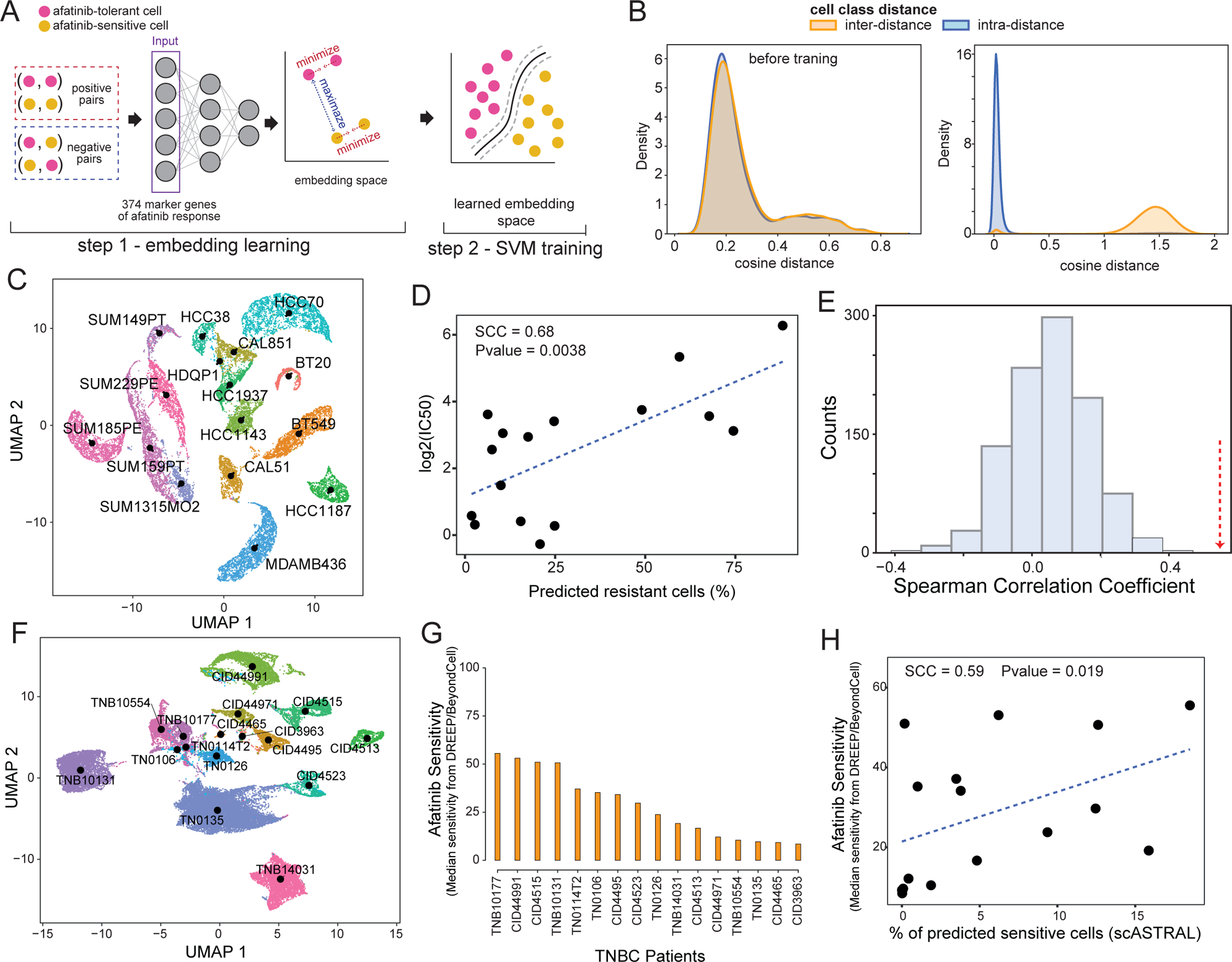
Single-cell afatinib response prediction in TNBC cells. (**A**) Schematics of scASTRAL. Contrastive learning is used in the first step to build an embedding space where positive examples (two cells of the same class) are separated from negative examples (two cells of different classes). In the second step an SVM classifier is trained on the learned embedded space to predict the class (i.e. afatinib-tolerant or afatinib-sensitive) of novel cells. (**B**) Inter- and intra-cosine distance between sensitive and tolerant MDAMB468 cells before and after the training. Intra-distance is the cosine distance among cells of the same afatinib response class, while inter-distance is the distance among cells belonging to two different afatinib response classes (**C**) UMAP representation of 22,724 triple negative breast cancer cells from 16 cell lines. (**D**) Spearman Correlation Coefficient (SCC) between scASTRAL predicted *resistance* to afatinib and experimentally estimated IC50. (**E**) Histogram of SCC values computed between scASTRAL predicted sensitivity to afatinib and experimentally estimated IC50 using 374 random genes. 1,000 simulations were performed. Red arrow indicates SCC value obtained using the 374 afatinib response marker genes we identified with our retrospective lineage tracing approach. (**F**) UMAP representation of 41,189 triple-negative breast cancer (TNBC) cells extracted from treatment-naïve primary tumours of 16 patients. (**G**) Estimation of the proportion of cells sensitive to afatinib using DREEP and BeyondCell tools. (**H**) Spearman Correlation Coefficient (SCC) between scASTRAL-predicted afatinib sensitivity and estimated sensitivity obtained from DREEP and BeyondCell tools.

We trained scASTRAL model on the labelled 1,541 MDAMB468 control cells (Methods) and then assessed its performance in predicting afatinib sensitivity on 22,724 single-cell transcriptional profiles from 16 TNBC cell lines (Figure 5C). Specifically, single cell data of 11 out of the 16 cell lines (15,022 cells) were obtained from the cell line breast cancer atlas we recently published (26), while data from the remaining 5 cell lines (7,702 cells), were de-novo sequenced using the DROP-seq platform (69) (Methods). We then employed scASTRAL to project the 22,724 triple negative cells into the learned afatinib-response space and classify each cell as either afatinib-sensitive or afatinib-resistant.

To evaluate scASTRAL’s performance in predicting afatinib sensitivity, we converted predictions to the cell-line level by computing the fraction of predicted afatinib-sensitive cells in each cell line and correlated these values with the experimentally determined afatinib IC50 values of each cell line. Afatinib IC50 values were retrieved from the Genomics of Drug Sensitivity in Cancer (GDSC) database v2 (34) or measured de-novo when unavailable (Supplementary Table 09). As shown in Figure 5D and Supplementary Table 10, scASTRAL’s predicted cell-line sensitivity exhibited a significant correlation with the corresponding experimentally determined IC50 values (SCC=0.68, P=0.0038). Consistent with expectations, our model’s predictive accuracy for resistance in treated cells significantly diminished. scASTRAL accurately classified only 5% of Day 3 cells, 12% of Day 6 cells, and a mere 2% of Day 9 cells as resistant. This decline in predictive performance is probably due to the disruption of marker gene expression induced by afatinib treatment. Indeed, as shown in Supplementary Figure 09, 230 out of the 374 marker genes consistently exhibited differential expression across all time points, indicating compelling evidence of treatment-induced alterations.

Next, to assess the predictive power of our 374 marker genes, we performed a series of Monte Carlo simulations (Methods) to evaluate the performance of randomly selected gene sets in predicting afatinib sensitivity. Specifically, in each simulation: (*i*) we randomly selected 347 genes from the overall gene pool; (*ii*) we trained scASTRAL on MDAMB468 control cells; (*iii*) we used the trained model to predict the afatinib response of the 16 TNBC cell lines described above; and (iv) we evaluated the performance of scASTRAL correlating predicted sensitivities with the experimentally estimated ones. We repeated this process 1,000 times. As shown in Figure 5E, our 374 marker genes consistently outperformed the randomly selected gene sets, suggesting that they possess superior predictive power for afatinib sensitivity. Next, we employed the Permutation Feature Importance (PFI) method (70) to evaluate the relative significance of the 374 marker genes associated with afatinib response that we had identified. The PFI algorithm is designed to focus solely on the predictive performance of the model, evaluating the importance of a feature by quantifying the increase in the model’s prediction error when the feature values are permuted (Methods). This analysis disclosed that out of the 374 genes, 212 had a positive PFI score (Supplementary Table 11), indicating their heightened significance in the model’s predictions. Interestingly, 41 of these significant genes, which constitute 19% of the total, are among those modulated by IGFBP2 overexpression.

Finally, to evaluate the applicability of scASTRAL on bulk transcriptomic profiles, we aggregated the single-cell expression profiles of the 16 TNBC cell lines (Figure 5A) into bulk profiles and applied scASTRAL to these aggregated data. This yielded estimates of the resistance probability for each cell line. We then mirrored our validation approach for single-cell data by correlating the scASTRAL-estimated probabilities with experimentally determined IC50 values for these cell lines. We observed a significant spearman correlation coefficient of 0.49 (Supplementary Figure 10) between the estimated probability of resistance and the IC50 values, indicating that scASTRAL may be readily adaptable for use with bulk transcriptomic data.

Cancer cell lines often exhibit limited transcriptomic heterogeneity, lacking the full complexity of actual tumors. Therefore, to demonstrate a possible clinical applicability of scASTRAL, we tested our methodology to a series of single-cell datasets derived from primary sites of individuals diagnosed with triple-negative breast cancer (TNBC). To this end, we assembled 41,189 single-cell transcriptional profiles encompassing 16 treatment-naive primary TNBC patients (71, 72) (Figure 5F). Since experimental data on afatinib sensitivity for these patients is unavailable, we employed an ensemble prediction approach using two state-of-the-art bioinformatics tools, DREEP (73) and Beyondcell (74), to estimate afatinib sensitivity from single-cell expression profiles (Figure 5G) (Methods). This strategy enabled us to estimate the proportion of afatinib-sensitive cells within each patient’s tumor cell population, which served as a reference for evaluating the performance of scASTRAL. As shown in Figure 5H, we observed a spearman correlation coefficient of 0.59 (p-value: 0.019), suggesting that our scASTRAL model can predict afatinib sensitivity with reasonable accuracy in TNBC patients as well. While this preliminary analysis provides some encouraging evidence for the potential clinical utility of scASTRAL, it is crucial to recognize that further validation studies involving larger datasets of patients treated with anti-EGFR therapy are essential to fully evaluate the model’s performance on more complex data as primary tumour’s biopsy. However, such data is currently unavailable.

Overall, these findings demonstrate the potential of our retrospective lineage tracing approach to identify biomarker genes predictive of drug response, which could potentially be utilized to stratify TNBC patients.

## Discussion

Owing to its inherent genetic complexity and the absence of recurrent oncogenic alterations, TNBC currently has limited treatment options beyond conventional chemotherapies (3, 5). Although the majority of primary TNBCs exhibit increased expression of EGFR due to an increased gene copy number, EGFR-targeted therapies have demonstrated variable and unpredictable clinical responses in TNBC. Hence, novel approaches are urgently needed to identify drug response biomarker genes that can effectively stratify TNBC patients for tailored EGFR-targeted therapies (18–22). Although clinical studies of EGFR inhibitors across various tumour types have not shown sufficiently high response rates to warrant their use in unselected patients, durable responses have been observed at low frequencies across several epithelial tumour types (75, 76). This highlights the need for novel approaches for robust biomarker identification strategies to accurately predict patient outcomes and optimize the use of EGFR-targeted therapies in TNBC.

In this study, we combined single-cell transcriptomics, cellular barcoding, and time-resolved computational analyses to provide a comprehensive transcriptional characterization of MDAMB468 cells in response to afatinib treatment, a potent TKI that irreversibly inhibits both EGFR and HER2 proteins.

Retrospective lineage tracing analysis uncovered a pre-existing subpopulation of MDAMB468 afatinib-tolerant cells exhibiting distinct biological features, such as elevated mRNA levels of the IGFBP2 gene. We provide experimental validation that IGFBP2 overexpression is sufficient to make TNBC cells tolerant to afatinib treatment through activation of the compensatory IGF1-R signalling pathway in MDAMB468 cells. Additionally, prospective lineage tracing analysis revealed the activation of several mechanisms that contribute to drug adaptation, including lysosome biogenesis, reactive oxygen species homeostasis, and fatty acid metabolism (48–54).

Our approach not only provided insights into the molecular mechanisms of afatinib resistance in MDAMB468 cells but also yielded a valuable tool for predicting drug response in other TNBC cells. Indeed, by leveraging deep learning techniques we devised an algorithm, scASTRAL, to accurately predict afatinib sensitivity from the expression levels of the 374 genes were identified through retrospective lineage tracing strategy. This signature demonstrated its generalizability by accurately predicting afatinib response in both publicly available and de novo single-cell datasets of TNBC cell lines and primary tumors. However, further studies are necessary to investigate whether the transcriptional programs linked to afatinib resistance are conserved across different tumor types and shared by other tyrosine kinase inhibitors (TKIs).

## Conclusions

Unraveling the distinct responses of cancer cell clones to treatment is crucial for understanding how intra-tumoral heterogeneity shapes drug effectiveness and identifying novel druggable targets for personalized cancer therapy. Our study highlights the importance of lineage tracing strategies, enabling the detection of subtle nuances that may go unnoticed by conventional clustering approaches reliant on gene expression alone. Our findings showcase the promising potential of lineage tracing techniques in reconstructing cancer clonal evolution under therapeutic interventions and demonstrate effectiveness in identifying predictive biomarker genes that can guide personalized treatment strategies for cancer patients.

## Methods

### Cell culture

HEK293T, MDAMB468, HCC1806 and HDQP1 were obtained from ATCC biobank. HEK293T and HDQP1 cells were cultured with DMEM supplemented with 10% foetal bovine serum (FBS), 1% L-glutamine, 1% penicillin/streptomycin (Euroclone, Milan, Italy), whereas MDAMB468 and HCC1806 cells were cultured with RPMI supplemented with 10% foetal bovine serum (FBS), 1% L-glutamine, 1% penicillin/streptomycin (Euroclone). Cells were kept at 37 °C under 5% CO_2_ atmosphere. SUM series cell lines were obtained from BioIVT biobank. SUM149PT, SUM185PE, SUM225CWN, SUM229PE and SUM159PT were cultured with Ham’s F12 medium (Gibco) supplemented with 5% heat-inactivated FBS (Gibco), 10 mM HEPES (Sigma-Aldrich), 1 μg/ml hydrocortisone (Sigma-Aldrich), 5 μg/ml insulin (Sigma-Aldrich) and 1% L-glutamine (Euroclone). SUM1315MO2 were cultured with the Ham’s F12 medium supplemented as described above with in addition 10 ng/mL EGF. No penicillin-streptomycin was added to the SUM cell line medium, as recommended by the suppliers.

### dCAS9 plasmids and sgRNA cloning

pRDKCE2B-EFS-dCas9-KRAB-2A-Blast, pRDVCE2B-EFS-dCas9-VPH-2A-Blast, pRSG16N-U6-sg-UbiC-TagRFP-2A-Neo and pRSG17H-U6-sg-UbiC-TagGFP2-2A-Hygro were obtained from Cellecta (#SVKRABC9E2B-PS, #SVVPHC9E2B-PS, #SVCRU616N-L and #SVCRU617H-L respectively). sgRNA used to overexpress or downregulate the IGFBP2 gene was designed using the CRISPick Broad Institute tool (https://portals.broadinstitute.org/gpp/public) (78, 79) (Supplementary Table 11). IGFBP2 protospacer sequences were synthesized, hybridized, phosphorylated, and inserted into pRSU6-gRNA plasmids using BbsI sites.

### Lentiviral packaging and transduction of dCAS9 plasmids

Lentiviral packaging was performed following the manufacturer’s instructions (Cellecta Mountain View, CA). Briefly, 24 hours before transfection, 4 x 10^6^ of HEK293T cells were plated in a 100-mm tissue-culture dish. On the day of transfection, 2 μg of the plasmid of interest was combined with 10 μg of the packaging plasmid mix (Cellecta, psPAX2: pMD2.G) in DMEM without serum or antibiotics in the presence of Plus Reagent™ and Lipofectamine™ (Life Technologies). Cells were incubated with the DNA/Plus Reagent™/Lipofectamine™ mix for 24 hours. Viral supernatant was collected 48 hours after transfection, filtered through a 0.45 μm PES filter and concentrated using LentiFuge™ Viral Concentration Reagent (Cellecta) following the manufacturer’s protocol. The concentrated lentiviral particles were re-suspended in PBS and stored at −80 °C. 5 x 10^5^ of MDAMB468 and HCC1806 cells were seeded in a 6-well plate and transduced with lentivirus to stably express dCas9-KRAB or dCas9-VHP in the presence of 8 μg/mL of LentiTrans™ Polybrene Transduction Reagent (Cellecta, #LTDR1) at a MOI of 0.5. 72 hours after transduction, 2.5 μg/mL of Blasticidin (Thermo Fisher Scientific, Waltham, MA, USA) was added to MDAMB468 cells, while 1 μg/mL of Blasticidin was used for the HCC1806 cell line. Once dCas9-KRAB and dCas9-VHP stable cell lines were obtained, they were transduced as previously with pRSU6-gRNA lentivirus to modulate IGFBP2 expression with a MOI of 0.5. After 5 days of selection, sgRNA RFP-positive or GFP-positive cells were further sorted using BD FACS ARIA III.

### HDQP1 cell transfection

The pCMV6-AC-IGFBP2-GFP expression vector encoding human IGFBP2 (NM_000597) fused to the GFP in the C-terminal region was purchased from Origene Technologies (#RG202573). This plasmid was used to obtain HDPQ1 cells stably overexpressing the IGFBP2 gene. Lipofectamine 2000 (Life Technologies, Grand Island, NY, USA) reagent was used to transfect HDQP1 cells according to the manufacturer’s instructions. The stable-transfected HDQ-P1 cells were selected in a medium containing 500 μg/mL G418 (Life Technologies). After 7 days of selection, cells were sorted with BD FACS ARIA III to select only the transfected population.

### CloneTracker XP barcode transduction

The MDAMB468 barcoded cell population was generated using CellTracker XP 10M Barcode-3’ Libraries (Cellecta, #BCXP10M3VP-V). Overall, 50,000 cells were transduced with the library with LentiTrans™ Polybrene Transduction Reagent at a low MOI (0.05) to yield only 5% infection. A population with 2,500 unique different barcodes was obtained. 72 hours post-infection, 1 μg/mL of Puromycin (Thermo Fisher Scientific) was added to the medium to select only transduced MDAMB468 cells. After 5 days, selected cells were checked by measuring the number of cells expressing Venus using the BD Accuri^TM^ C6 flowcytometer (BD Biosciences, Franklin Lakes, NJ, USA). The cells were 100% positive and thus expanded in cultured medium with 1 μg/mL of Puromycin.

### Time series experiment

5 x 10^5^ barcoded cells were plated in triplicate in a 6-well plate and allowed to adhere for 24 hours. Control cells were also plated in triplicate and viability checked over the course of the experiment. Cells were then treated with incremental concentrations of afatinib (Selleckchem cat. Num. S1011) starting from 250 nM with each treatment lasting 72 hours. Afatinib concentration was doubled every nine days up to 2 μM. Every 3 days, cell viability was measured for both treated and untreated cells and 1 x 10^4^ of treated cells were collected and cryopreserved from each treated replicate as follows. Cells were centrifuged at 1,100 rpm for 3 min and then resuspended with resuspension buffer (RB) (100% of FBS) to obtain a final concentration of 1,000 cells/µl. Then, 10 µl of this cell suspension was added to 90 µl of RB to obtain a final concentration of 100 cells/µl (total: 1 x 10^4^ cells). Finally, 100 µl of cold freezing buffer (FB) (100% of FBS and 5% DMSO) was added before placing the cells at −80°C. We chose to initiate afatinib treatment at 250 nM based on the lowest reported sensitivity of a breast cancer cell line to afatinib from GDSC1 screening, available at https://www.cancerrxgene.org. The chosen concentration range intentionally mimics a drug adaptation process, ranging from the minimal level at which a breast cancer cell line responds to an EGFR inhibitor to the highest concentration, which is twice the IC50 value of Afatinib in the MDA-MB-468 cell line. Regarding the 9-day period between doubling the drug concentration, this was determined after extensive experimentation. This interval provided sufficient time for the cells to adapt to the drug treatment and maintain survival. When we attempted to double the drug concentration more frequently, we encountered significant variability in cell viability and experiment reproducibility. The current strategy, with a 9-day interval, ensured consistent results throughout the experiments.

### Single cell RNA-sequencing of MDAMB468 cells with 10x chromium platform

The cryopreserved samples were thawed in a 37 °C water bath and washed multiple times with MACS® Separation Buffer (Miltenyi biotech). Cell viability of the single cell suspension was measured with Trypan Blue using a LUNA-II™ Automated Cell Counter (Logos Biosystems). Cells were then filtered through a 40-μm filter, centrifuged at 300×g for 5 min and resuspended to obtain a final concentration of 1 x 10^3^ cells/ µl. The cell suspensions were then processed to generate single-cell libraries using a Chromium Single Cell Gene Expression 3′ Library and Gel Bead kit v3.1 following the manufacturer’s instructions (10x Genomics). Briefly, the cells were suspended in reverse transcription reagents and injected into microfluidic chips of the 10x Chromium instrument, along with gel beads, and segregated in a nanoliter-scale Gel Beads-in-Emulsion (GEMs). Reverse transcription was carried out on the GEMs in a MiniAmp Thermal Cycler (Thermo Fisher Scientific) with the following protocol: 53 °C for 45 min, 85 °C for 5 min, and hold at 4 °C. After, reverse-transcribed samples were purified with the Recovery agent and isolated with the Dynabeads Cleanup Mix. The cDNA was amplified in a Thermo cycler using the following protocol: 98 °C for 3 min, 12 cycles of (98 °C for 15 sec, 63 °C for 20 sec, 72 °C for 1 min), 72°C for 1 min, and hold at 4°C. The quality of the amplified cDNA was quantified with the BioAnalyzer High Sensitivity DNA kit D5000 (Agilent Technologies) and its concentration was measured using the Qubit Fluorometer. Finally, amplified cDNAs were fragmented, end-repaired and A-tailed with SPRIselect Reagent Kit (Beckman Coulter). The post-ligation products were amplified using the following protocol: 98 °C for 45 sec, 10 cycles of (98 °C for 20 sec, 54 °C for 30 sec, 72 °C for 20 sec), 72 °C for 1 min and hold at 4 °C. The sequencing-ready libraries were purified with SPRIselect and the quality were quantified with the BioAnalyzer High Sensitivity DNA kit D1000 and the concentrations were measured using the Qubit Fluorometer. NovaSeq 6000 SP 100 cycles flow cell was used to sequence the libraries.

### Specific CloneTracker XP barcode single cell library

To optimize the capture of the CloneTracker XP barcode at the single cell level, a parallel single cell library preparation was performed from the amplified cDNAs by separately amplifying the CloneTracker XP barcode amplicon with a custom P7 primer that targeted it (Supplementary Table 12). CloneTracker XP barcode was amplified according to the following protocol: 98 °C for 45 sec, 14 cycles of (98 °C for 20 sec, 54 °C for 30 sec, 72 °C for 20 sec), 72 °C for 1 min. PCR products were then SPRI-purified and the quality were quantified with the BioAnalyzer High Sensitivity DNA kit D1000 and the concentrations were measured using the Qubit Fluorometer. Single cell CloneTracker XP barcode libraries were sequenced as spike-ins alongside the parent RNA-seq libraries. This strategy to use a custom P7 primer with unique i7 index had the advantage to allow to separately analyse the barcoded scRNA-derived cDNA in the NGS data.

### CloneTracker XP barcode retrieval from scRNA-Seq reads

As shown in Figure 01C, the 48-nucleotide expressed barcode cassette of the Cellecta CloneTracker XP barcode library has a composite structure built by juxtapositioning 100 different 14-nt sequences (a.k.a. bc14) and 100,000 30-nt sequences (a.k.a. bc30) separated by a 4-nt (TGGT) anchor. Cellecta also provides the whitelist containing the 100 possible bc14, and 100,000 bc30 can appear at both sides of the anchor. Lineage barcodes were identified from the FASTQ file of read two, while cell barcode from read one of both the specific CloneTracker XP barcode single cell library and standard single cell library we produced for each sequenced time point. Specifically, for each fragment present in read two, the occurrence of the TGGT anchor was first searched allowing one possible mismatch. Then, if some correspondence was found, the 14-nt bases before (*i.e.* putative bc14) and the 30-nt bases after (*<u>i.e.</u>* putative bc30) the anchor sequence were extracted. Next, if putative bc14 and bc30 were in the whitelist provided by Cellecta, the lineage barcode represented by the string “bc14:bc30” was assigned to the corresponding cell retrieved from read one. On the other hand, if one of the putative lineage barcodes was not found in the Cellecta whitelist, we corrected it with the barcode in the whitelist having a Hamming distance equal to one, but only if there were no other barcodes in the whitelist with a Hamming distance of one. Finally, if both the putative bc14 and bc30 were not found in the Cellecta whitelist, the bc14 barcode sequence is corrected first as above and we corrected the corresponding bc30 only if bc14 correction was successful. This strategy allowed us to reduce the high computing demand required by correcting lineage barcodes for possible sequencing errors but without losing sensitivity. At end of this iterative process, each sequenced cell is associated with the most abundant pair of lineage barcodes bc14:bc30 we have found. Only reads where both bc14 and bc30, after correction, matched with the ones provided by Cellecta’s whitelist were used. This was implemented in an R script that made use of the Biostrings R package to perform string matching and a custom C++ function to compute the Hamming distances between two DNA sequences of the same length.

### MDAMB468 scRNA-seq reads alignment and expression quantification

Following demultiplexing, raw sequencing reads were aligned to the Hg38 human reference genome using the Cell Ranger tool version 6.1.2. For reads alignment, the GENECODE annotation v32 of Hg38 was employed, retaining only the genes with biotypes protein coding, lincRNA, and antisense. After reads alignment, cells with fewer than 5,000 UMI were discarded. Next, putative cell doublets and cells expressing a high fraction of mitochondrial reads were removed using the *filterCell* function from the gficf version 2 R package (27, 80, 81) available at https://github.com/gambalab/gficf. Specifically, the *filterCell* function employs loess regression to fit the relationship between the total UMI count in a cell (in log scale) and the ratio between the total UMIs falling in mitochondrial genes over the total UMI count. Cells in which the ratio deviates significantly from the expected value with an adjusted p-value (FDR) < 0.1 are discarded. To eliminate putative cell doublets, the same strategy is applied, but in this case, loess regression is used to fit the relationship between the total UMI count in a cell (in log scale) and the ratio between the number of detected genes over the total UMI count. Again, cells for which the ratio deviates significantly from the expected value with an FDR < 0.1 are discarded. According to our long experience in tha analysis of single cell data, this latter strategy is particularly useful for removing not only cell doublets but also cells that, for technical reasons, exhibit a high total UMI count concentrated in a small number of genes. The resulting highly covered 4,101 cells after these filtering processes were used for all downstream analysis. Furthermore, since cells of day 0 were sequenced using two distinct captures in two different flow cells, the possible batch effect was corrected using the *sva* function of CombatSeq R package (82). Finally, genes expressed in less than 50 cells and in less than 5% of sequenced cells were excluded only if their average expression, was less than 1.12 UMI. This approach is similar to that employed in (83).

### Bulk estimation of the number of lineages in MDAMB468 cells

Bulk estimation of the number of CloneTracker XP barcodes present in parental and ATPC MDAMB468 barcoded cells was performed with QuantSeq-Flex (Lexogen) technology as follows. First, total RNA was isolated from three independent biological replicates using the Qiagen RNeasy Mini kit (Qiagen), according to the manufacturer’s instructions. Then, library preparation was performed according to the manufacture’s protocol (Lexogene). The target-specific reverse transcription primers to specifically capture the Cellecta barcode were designed according to the guidelines outlined in the QuantSeq-Flex manual. All primers were composed of a partial Illumina P7 adapter extension followed by the target-specific sequence. A pool of 4 primers was used at 50 nM. Primer lengths were in the range of 44–50 nucleotides, as requested by the manufacture’s protocol. The target-specific first strand synthesis primers can be found in Supplementary Table 11.

### Cell viability assay

Cells were seeded in either a 96-well plate (4 x 10^4^ cell/well) or a 384-well plate (1 x 10^3^ cell/well). After overnight incubation at 37 °C, cells were treated either with drugs, at the selected dilutions, or with DMSO as negative control (in technical triplicate) and incubated at 37 °C for 72 hr. Then, cell viability was evaluated by measuring either luminescence or absorbance (490 nm) with the CellTiter-Glo® luminescent cell viability assay (Promega) or the CellTiter (Promega), respectively, using the GloMax**®** Discover instrument (Promega) according to the manufacturers’ protocol. Background luminescence or absorbance values were measured in wells without cells and with only culture medium. Background values were subtracted from experimental values. All drugs used in this study were purchased from Selleckchem.

### Drug combination assay

4 x 10^4^ of MDAMB468 cells were seeded in a 96-well plate and incubated overnight at 37 °C. Afatinib and GSK-1904529A drugs were prepared in different dilutions and then combined in all possible drug pairs to generate a 5 x 5 or 5 x 6 drug combination matrix. Then, cells were exposed to either single agent drugs or to the drug pairs of the drug combination matrix, while negative controls were treated with DMSO (each treatment was performed in triplicate). Following 72 hr incubation at 37 °C, cell viability was measured with the CellTiter (Promega), and the absorbance was read at 490 nm with the plate reader GloMax® Discover instrument. The Afatinib-GSK-1904529A drug interactions and expected drug responses were calculated with the Combenefit tool (84) using the Loewe additivity model.

### Generation of spheroids and image acquisition

Cells grown as a monolayer were detached to generate a single-cell suspension that was then diluted to 2.5 × 10^4^ (for 5,000 cells per spheroid) cells per millilitre of ice-cold medium. The Cell Basement Membrane (ATCC, #ATCC-ACS-3035) was thawed on ice overnight and added at a final concentration of 2.5% with ice-cold pipette tips to the cell suspension. A volume of 200 µl of this cell suspension was added to each well of an Ultra-Low Attachment (ULA) 96-well plate with a round or conical bottom (PerkinElmer, #SPA6055330). The spheroid formation was initiated by centrifugation of the plates at 300g for 10 min. The plates were incubated under standard cell culture conditions at 37 °C, 7% CO_2_ in humidified incubators. When we studied afatinib response, the drug was added as follows. After 24 hours, when spheroids were formed, 100 μl of medium was removed and the treatment was performed adding 100 μl of medium with 2 μM of afatinib into each well. Treatments were renewed after 3 days in new fresh complete medium. Pictures were taken before adding treatments and then at day 3, 5 and 7 after treatment. Images were acquired with the High Content Analysis System *Operetta CLS* (PerkinElmer) with temperature (37 °C) and CO_2_ (5%) control. They were acquired in several planes and the area analyses were performed on maximum projections by Harmony software (PerkinElmer). Volumetric analyses were performed by stack processing with 3d analysis.

### RNA extraction, reverse transcription, and quantitative RT-PCR (qPCR)

Total RNA from cell lines was extracted usingthe Qiagen RNeasy Mini kit (Qiagen, Hilden, Germany) following the manufacturer’s instructions. A total of 1 μg of total RNA from each sample was used to obtain double strand cDNA with the QuantiTect Reverse Transcription Kit (Qiagen). Quantitative Real-Time PCR (qRT-PCR) was performed and for each PCR reaction, 10 μL of 2× Sybr Green (LightCycler 96, Roche Molecular Systems, Inc.), 200 nM of each primer, and 20 ng of the cDNA, previously generated, were used. Relative gene expression was determined using comparative C(T) method, as described elsewhere. RP18S was used as a housekeeping gene.

### Clonogenic survival assay

5 x 10^4^ cells were plated into a 12-well plate and allowed to adhere for 24 hours. Then the cells were treated with different concentrations of Afatinib and incubated for 10 (Figure 3C) or 5 days (Figure 3E). Fresh media was added on the fifth day. On the tenth day, the media was removed from the dishes and washed once with ice-cold PBS. The colonies were fixed and stained with a solution containing crystal violet for 45 minutes on a rocking platform. The dishes were rinsed three times with PBS and air-dried. Then, the stained crystal violet was resolved with PBS-0.1% SDS and absorbance at 590 nm was determined. The data were performed in triplicate and are shown as mean ± SD.

### QuantSeq 3′ mRNA-Sequencing and bioinformatics analysis

Total RNA was purified from three biological replicates by Qiagen RNeasy Mini kit (Qiagen), according to the manufacturer’s instructions. Total RNA was quantified using the Qubit 4.0 fluorometric Assay (Thermo Fisher Scientific). Libraries were prepared from 125 ng of total RNA using the NEGEDIA Digital mRNA-seq research grade sequencing service (Next Generation Diagnostic srl) which included library preparation, quality assessment and sequencing on a NovaSeq 6000 sequencing system using a single-end, 100 cycle strategy (Illumina Inc.). The raw data were analysed as follow. Briefly, Illumina novaSeq base call (BCL) files were transformed into fastq files through bcl2fastq (version v2.20.0.422, Illumina Inc.), and sequence reads were trimmed using bbduk software (bbmap suite 37.31, Joint Genome Institute, Walnut Creek, CA, USA). Alignment was performed on hg38 reference assembly (Ensembl Assembly 93) with star 2.6.0a (GPL v3, open source) and gene expression levels were determined with htseq-count 0.9.1. Gene expression normalization and differentially expressed genes were identified using edger package (85) in the R statistical environment. For differential expression analysis, only genes with an average CPM more than 5 in at least one of the two conditions were considered.

### Differential expression analysis and clustering analysis of single cell data

Differentially expressed genes in Figure 2C between tolerant and sensitive afatinib cells at time 0 (*i.e.* untreated cells) were identified using the *FindMarkers* function implemented in Seurat R package v3 with logfc.threshold and min.pct parameters set to 0. All clustering analyses were performed with Seurat.

### Pseudotime analysis of tolerant clones and time-course differential expression analysis between cells of the dominant and neutral clones

The 2,836 single cell expression profiles associated with afatinib-tolerant clones were divided into two groups according to the behaviour of their clone of origin, *i.e.* “dominant” or “neutral”. The expression matrix of dominant clones comprised 1,293 cells while the matrix of neutral clones comprised 1,543 cells. Then *psupertime* function of psupertime R package (45) was used to infer the pseudotime order of cells in each independent matrix. Psupertime R function was run with default parameters and using as cell labels their sequencing day (i.e., 0, 3, 6, 9 days). Next, for both matrices the expression profile of each gene was (*i*) ordered according to the reconstructed pseudotime; (*ii*) smoothed using the moving median method and (*iii*) finally divided into 50 expression bins according to the inferred pseudotime. To smooth gene expression with the moving median method, we used *rollmedian* function of the zoo R package with a rolling window equal to 51. After these three steps the expression profiles of each gene in both matrices was represented by 50 pseudotime points. Finally, to identify upregulated genes in the cells associated with dominant clones across the first 9 days of treatment we used the splineTimeR package (46). With this tool we compared the reconstructed time-course profile of a gene across cells of the dominant clones with its reconstructed time-course profile across cells of the neutral clones. Up-regulated genes were defined as the genes with an FDR < 5% and a positive average fold change across the 50 binned pseudotime points.

### Estimation of EGFR copy number alterations, mutations, and expression in TNBC patients

Two independent cohorts of TNBC patients were used to estimate gene copy number alterations, mutations and expression reported in Supplementary Figure 01. The first cohort composed of 192 TNBC patients was obtained from the Genomic Data Common (GDC) portal (86). GDC raw bulk expression, mutational data and copy number variation data along with clinical information were retrieved using TCGAbiolinks R package v.2.25.3 (87). The second cohort was composed of 465 TNBC Asiatic patients (5) for whom gene expression, gene copy number alteration and mutations were available was gathered from the NODE database after the author’s authorization. The raw expression counts downloaded from GDC were first normalized with edgeR package (85) and transformed in log2(CPM + 1), while those from the Asiatic cohort were already normalized as FPKM values, and thus we only transformed them into log2(FPKM + 1). A sample in the GDC cohort was defined to have EGFR copy gain if the number of copies was 3 or 4, while EGFR amplification was defined when this number was greater than 5. For the Asiatic cohort, EGFR gain and amplification were defined based on GISTIC scores, respectively, +1 and +2. For both cohorts, mutations were considered only if they were in the coding region (i.e., missense, non-sense, in-frame and out-of-frame deletions). Next, the deleterious tag was assigned when at least two functional annotations among VEP, PolyPhen and SIFT reported a negative impact of the mutation on protein function. Finally, EGFR activating mutations tag was conferred based on indications in (88).

### Cell migration assay

Transwell chambers (8-μm pores) (Euroclone) were used to perform transwell migration assays. Briefly, breast cancer cells were plated in the upper side of the transwell chamber in a serum-deprived medium. Next, as chemoattractant, 300 μl of complete medium was added in the lower chamber. After 24 hours, cells migrated to the lower side of the chambers were stained with crystal violet solution (crystal violet 0.05%, methanol 20%). Then, crystal violet in the chamber was de-stained with PBS-0.1% SDS solution and read at 570 nm.

### scRNA library preparation of Drop-SEQ and bioinformatics analysis

Single cell transcriptomics of the SUM149PT, SUM185PE, SUM22PE, SUM159PT and SUM1315MO2 cell lines was performed with DROP-seq technology (69) and library preparation as described in Gambardella et al. (26). scRNA libraries were sequenced with NovaSeq 6000 machine using an SP 100 cycles flow cell. Raw reads pre-processing was performed as described in (81). Only high depth cells with at least 5,000 UMI were retained and used to test the scATRAL tool. The alignment pipeline can be found at https://github.com/gambalab/dropseq.

### Gaussian processes to estimate significance in dose-response curves

We used Gaussian processes to evaluate the significance of dose-response curve trend differences. Gaussian processes computation was performed as in (89). Briefly, we first reconstructed for each condition the dose response trend by interpolating the response values at each tested concentration and then we identified differences in the two trends under consideration in terms of log-likelihood, by computing the related Bayesian factor.

### Estimation of the cellular frequency over time of the 192 afatinib-tolerant clones

To compute cellular frequencies of an afatinib-tolerant lineage over time, we estimated the percentage of cells associated with it in each sequenced time point. For each time-point, this percentage was simply estimated as the number of cells associated with a specific lineage divided by the total number of sequenced cells.

### scASTRAL: architecture

Contrastive learning is a machine learning approach that aims to create an embedding space where positive examples (*i.e.* two cells with the same label) are separated from negative examples (*i.e.* two cells with different labels). In our case, a contrastive autoencoder was used as a model of scASTRAL. The encoder consisted of an input layer composed of 374 neurons, *i.e.* the number of afatinib response biomarker genes identified with their retrospective lineage tracing strategy. The input layer was followed by two hidden layers comprising 64 and 32 neurons, respectively. A ReLU (Rectified Linear Unit) activation function was used. The decoder was symmetrical to the encoder.

### scASTRAL: training

We used the 1,541 MDAMB468 control cells labelled as afatinib-tolerant or afatinib-sensitive as a training set. Before starting the training, the matrix of normalized CPM counts was cut to the 374 afatinib response biomarker genes we identified in Figure 3C and the expression of these genes rescaled using tf-idf transformation (80). This dataset was then randomly split into 5 batches of equal size by ensuring that the number of afatinib-tolerant and afatinib-sensitive cells were similar across the different batches. Next, a contrastive model was trained on each batch as follows. First, a positive and a negative example was randomly built for each cell of the batch. This means that for each batch the number of examples on which training is performed was always equal to the number of cells in the batch multiplied by two. Second, the contrastive model was trained using a loss function composed by the weighted sum of three terms: the cosine embedding loss, the reconstruction error, and the latent space regularization factor. Specifically, the cosine embedding loss maximizes the cosine distance between pairs of cells labelled with −1 (*i.e.* negative example) while maximizing the cosine similarity between pairs of cells labelled with +1. (*i.e.* positive example). At the same time, the reconstruction error assures that the decoder was capable of reconstructing original data from the latent representation generated by the encoder. This allows the model to also integrate in the final embedding space the dependencies among the 374 genes we used as input. Third, an SVM classifier is trained using the reconstructed embedding space of the considered batch of cells. Specifically, the SVM classifier was trained with a cosine kernel and with a regularization hyperparameter set to 100. Classification accuracy of the trained SVM of a batch was finally evaluated using the left-out cells. We used the Adam optimizer (90) with a learning rate of 0.0001 while the training was performed using an early stopping criterion, although the maximum number of epochs was set to 250. The metrics considered to compute improvements across epochs were the validation loss and the SVM classification accuracy. The training stopped if no improvements were obtained considering the last 20 epochs. The contrastive model hyperparameters were found by a grid search. Once the 5 contrastive models with associated SVM models were trained, the model with the highest classification accuracy was used to predict afatinib sensitivity at the single cell level as described below. scASTRAL was implemented in Phyton using PyTorch version 1.13.1. Code is available at the following address https://github.com/gambalab/scASTRAL.

### scASTRAL performance evaluation on single cells of TNBC cell lines

To validate scASTRAL and test its performance in predicting afatinib sensitivity of triple negative breast cancer cells, we used 22,724 single-cell transcriptional profiles from 16 TNBC cell lines (Figure 5D). Of these, 11 were obtained from Gambardella et al. (26), while five were de-novo sequenced with the drop-seq platform. Before being fed into the scASTRAL method, the raw UMI count matrix of each cell line was first normalized using sklearn to obtain CPM and then cut on the 374 afatinib response biomarker genes identified in Figure 3C. Next, the expression values of these genes were rescaled using tf-idf transformation (80) before applying scASTRAL. Finally, a cell was deemed to be tolerant or sensitive to afatinib if the SVM classification probability was greater than 0.75 otherwise it was considered undetermined. To convert predictions from the single-cell level to the cell-line level, we computed the fraction of predicted afatinib-tolerant cells in each cell line as the number of cells predicted to be tolerant divided by the total number of cells.

### scASTRAL performance evaluation on pseudobulk of TNBC cell lines

Pseudobulk profiles of each cell line were computed by summing the UMI counts of each gene across the sequenced cells of the corresponding cell lines. The corresponding count matrix was cut on the 374 marker genes of afatinib response and then fed to scASTRAL.

### scASTRAL performance evaluation on single cells of TNBC patients

To evaluate the performance of scASTRAL on single-cell data from TNBC patients, we compiled 41,189 single-cell transcriptional profiles from 16 treatment-naive primary TNBC patients from the studies conducted by Pal et al. (71) and Wu et al. (72). As experimental data on the afatinib sensitivity of these patients was not available, we reconstructed this information using an ensemble prediction generated by the DREEP (https://github.com/gambalab/DREEP) and Beyondcell (https://github.com/cnio-bu/beyondcell) tools. These methods employ unique drug signatures to assess the impact of a drug and have been demonstrated to effectively predict drug response at the single-cell level. DREEP possesses five distinct signatures for Afatinib, while Beyondcell has three. To estimate the proportion of afatinib-sensitive cells in each patient’s tumor cell population, we applied both tools to each patient’s cells and assigned the median values across the eight potential signatures as the percentage of sensitive cells for that patient. Consequently, we could estimate the percentage of afatinib-sensitive cells in each patient’s tumor cell population, which we then used as a benchmark to compare the performance of scASTRAL. Both tools were run using default parameters. Subsequently, scASTRAL was applied to single-cell data of the 16 TNBC patients, employing the same approach we used for single cell data of TNBC cell lines. Hence, before being fed into the scASTRAL tool, the raw UMI count matrix of each patient was normalized using sklearn to obtain CPM and then cut on the 374 afatinib response biomarker genes. Subsequently, the expression values of these genes were rescaled using tf-idf transformation before applying scASTRAL. Finally, a cell was considered tolerant or sensitive to afatinib if the SVM classification probability was greater than 0.75; otherwise, it was deemed undetermined. To convert predictions from the single-cell level to the patient level, we calculated the fraction of predicted afatinib-sensitive cells in patient as the number of cells predicted to be sensitive divided by the total number of cells.

### Permutation Feature Importance (PFI) Analysis

To assess the significance of features within our classification model, we utilized the permutation feature importance (PFI) algorithm (70), a versatile model inspection technique applicable to any estimator working with tabular data. This technique exclusively relies on input data and output predictions, enabling its seamless integration into the comprehensive end-to-end scAstral pipeline. The PFI of a feature is quantified as the decrease in a scoring metric when the values of that feature are randomly shuffled within a batch of data. In our case, we employed the ROC AUC as the scoring metric. This process is repeated 1,000 times using different seeds for the shuffling, and the mean score across repetitions is calculated, providing a robust metric for evaluating the importance of each feature.

### EGFR copy number estimation and afatinib response of TNBC cell lines

Gene copy number data for EGFR was extracted from the DepMap database (version 23Q4) and transformed into actual copy counts. TNBC cell lines were classified into three groups based on their EGFR copy number distribution: Neutral (2 copies), Gained (3-4 copies), Amplified (≥5 copies) and Deleted (<2 copies). Afatinib response data was acquired from the GDSC database, comprising both GDSC1 and GDSC2 datasets.

## Code availability

scASTRAL code is available at https://github.com/gambalab/scASTRAL. DropSeq Alignment pipeline can be found at https://github.com/gambalab/dropseq.

## Acknowledgments

We thank the TIGEM high content screening facility for helping in 3D-spheroid analysis. We thank the TIGEM bioinformatics core for helping in dose response curve analyses. We express our gratitude to Dr. Cathal Wilson for meticulously proofreading our manuscript. His invaluable efforts have significantly contributed to enhance the clarity and accuracy of our work.

## Author Contribution

SP performed time series experiment, 10x sequencing and all the experimental validations. MF performed computational analyses. GV setup DropSeq platform and performed sequencing. RA developed the drug prediction method. GG supervised the work, wrote the manuscript, and conceived the original idea.

