## Supplementary Figures for "Single cell lineage tracing reveals clonal dynamics of anti-EGFR therapy resistance in triple negative breast cancer"

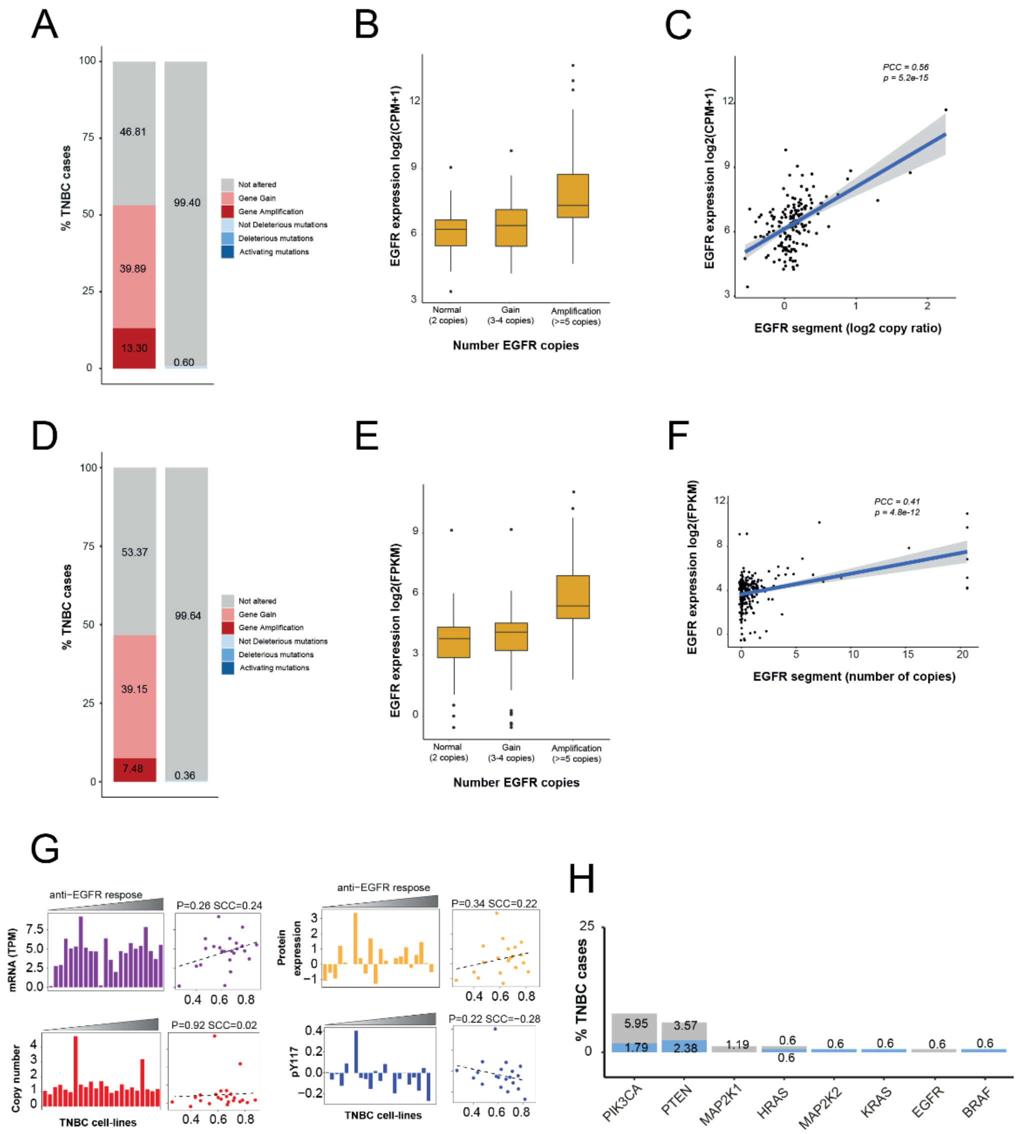

**Supplementary Figure 01 – EGFR genetic alterations in patients' cohorts. (A)** Bar plot reporting the percentage of TNBC patients from TCGA cohort with copy number alterations (left bar) or mutations (right bar) in EGFR gene. Specifically, copy number alterations were divided in gain (3-4 copies, pink) or amplification (≥ 5 copies, red), while mutations were separated in activating (dark blue), deleterious (light blue) and not deleterious ones (cyan). **(B)** EGFR gene expression distribution as a function of its copy number status in TNBC patients from TCGA cohort. **(C)** Correlation between EGFR gene expression and its copy number alteration status in TNBC patients from TCGA. EGFR copy number status is represented by the $\log_2$ copy ratio of its segment (x-axis). **(D)** Bar plot reporting EGFR genetic alterations at the copy number and mutational level (Asian cohort of patients). **(E)** EGFR gene expression as a function of its copy number status in TNBC patients from Asian cohort. **(F)** Correlation between EGFR gene expression and its copy number alteration status in TNBC patients from Asian cohort. EGFR copy number status is represented by the number of copies of the segment where it is located (x-axis). **(G)** Correlation between efficacy of anti-EGFR and EGFR status (mRNA, protein expression, copy number and EGFR phosphorylation status) in TNBC cell-lines. Data were downloaded from GDSC (Genomic of Drugs Sensitivity in Cancer) portal. Efficacy of anti-EGFR was computed as average $IC_{50}$ of all drugs targeting EGFR. **(H)** Bar plot showing the percentage of TNBC patients (y-axis) harboring both deleterious (blue) and not deleterious (grey) mutations in known anti-EGFRs resistance genes reported on x-axis.

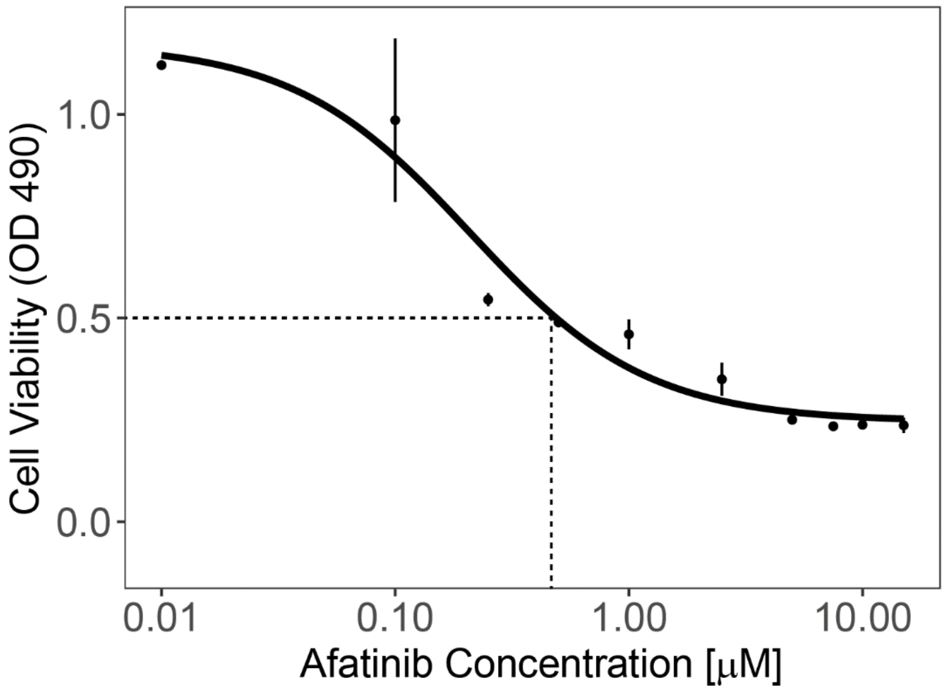


**Supplementary Figure 02 – MDA-MB-468 cell line responsiveness to Afatinib.** Dose-response curve in terms of cell viability (y-axis) after Afatinib treatment at the indicated concentrations (x-axis) on MDA-MB-468 cell line.

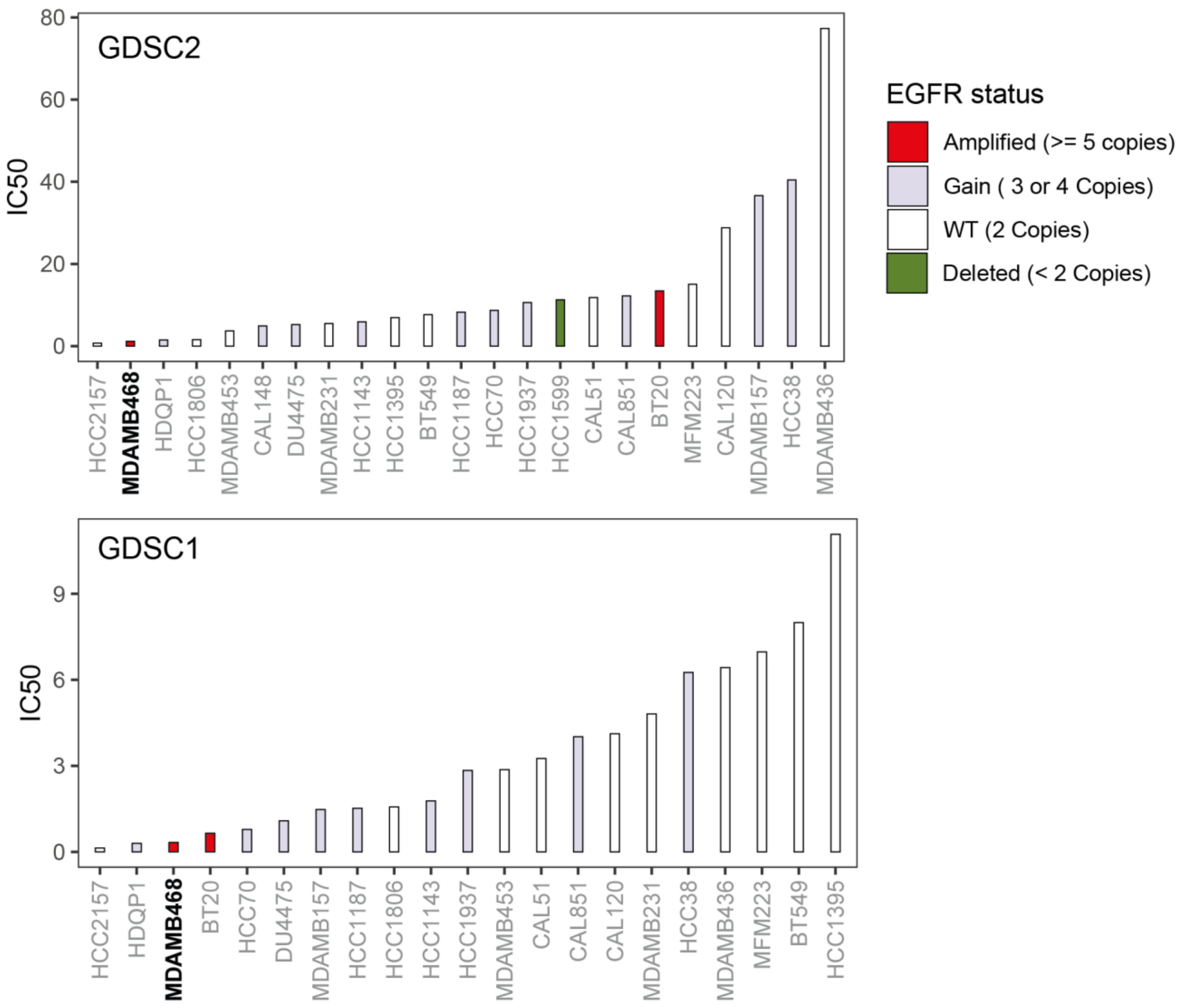


**Supplementary Figure 03 – Afatinib response of TNBC cell lines.** The afatinib responses were obtained from the Genomics of Drug Sensitivity in Cancer (GDSC) database (https://www.cancerrxgene.org). The cell lines are color-coded according to their EGFR copy number (Methods).

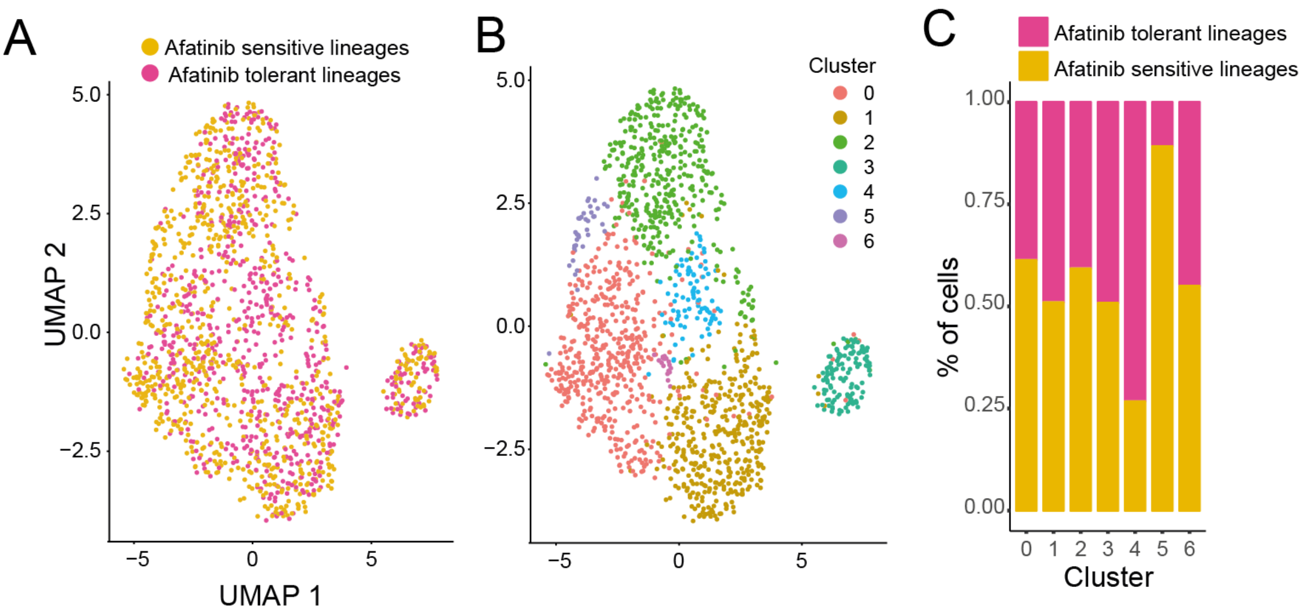


**Supplementary Figure 04 – Clustering analysis of Day 0 (untreated) cells. (A)** Uniform Manifold Approximation and Projection (UMAP) visualization of Day 0 cells. Cells are color-coded according to their lineage of origin, either afatinib-sensitive or afatinib-tolerant.. (**B**) Same as (A) where cells are color-coded according to their assigned cluster. (**C**) Proportion of afatinib-tolerant and afatinib-sensitive cells within each cluster.

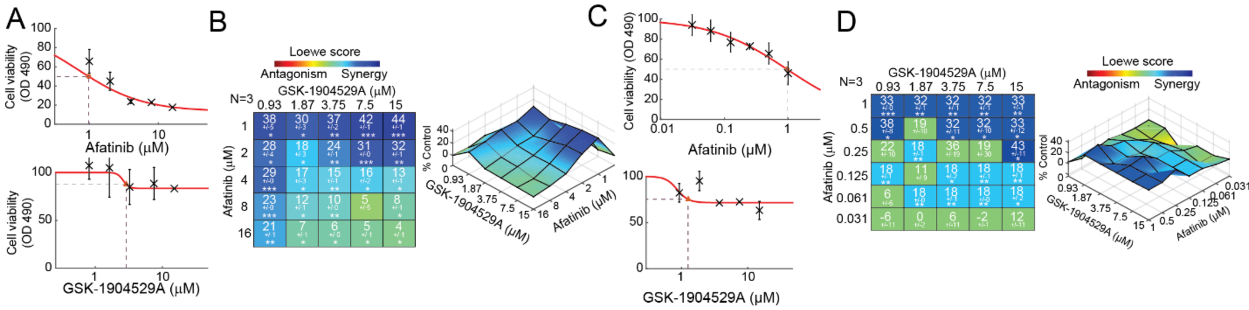


**Supplementary Figure 05 – Compensatory activation of IGF1-R and INSR pathways to overcome EGFR inhibition. (A)** Single agent dose-response data and fitted curve for Afatinib (upper) and GSK-1904529A (lower) on MDA-MB-468 parental cells. **(B)** (Left plot). Synergy scores calculated using Combenefit tool according to the Loewe additivity model (Methods) for different concentration of Afatinib and GSK-1904529A on MDA-MB-468 parental cells. The larger the value stronger is the synergism while negative values mean antagonism between the two drugs. The number below the synergy score is standard deviation. Asterisks below standard deviation indicate results that are statistically significant by the one-sample t-test (see methods). The degree of significance are as follows: * p < 5x10-2; ** p<10-3; *** p<10-4. The number of biological replicates (N) is indicated at the top left of the matrix. (Right plot) Combination dose-response surface, expressed as a percentage of the control value. Overlain on the dose response surface are the Loewe synergy scores. **(C)** Same as (A) but where concentration of Afatinib vary from 31nM to 1µM. **(D)** Same of (B) but where concentration of Afatinib vary from 31nM to 1µM.

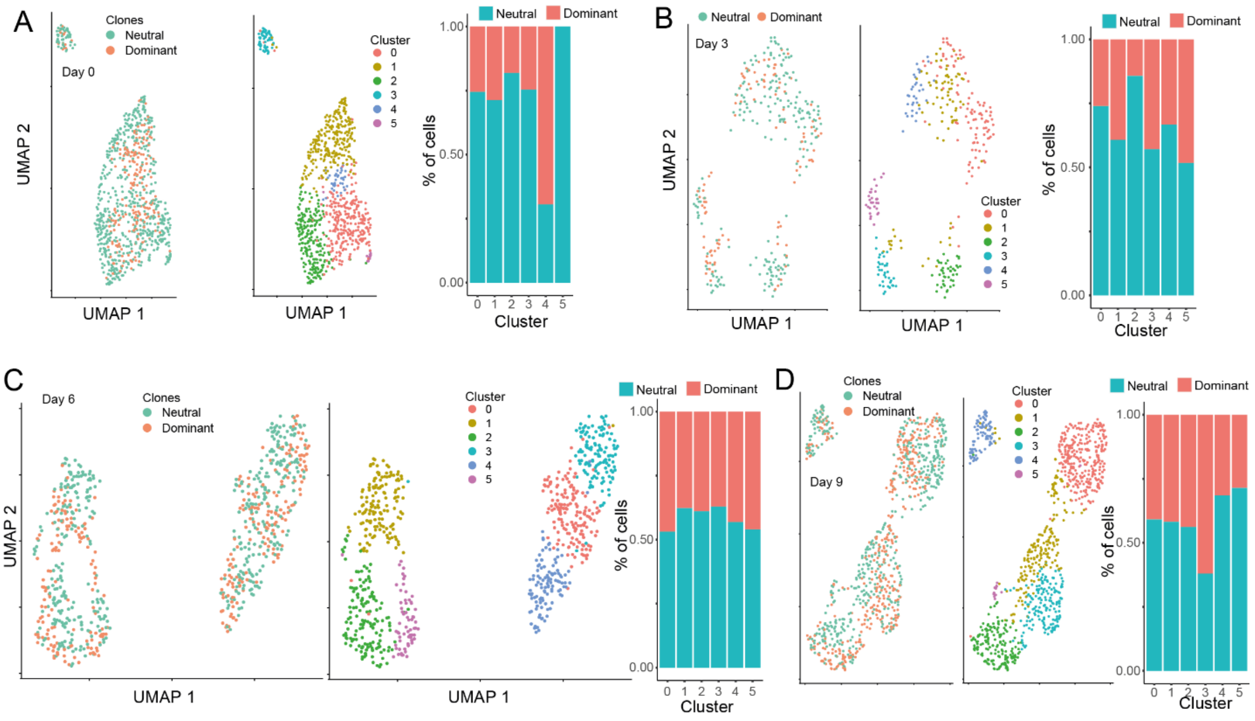


**Supplementary Figure 06 – Clustering analysis of afatinib tolerant cells. (A)** Left panel: Uniform Manifold Approximation and Projection (UMAP) visualization of Day 0 cells where cells are color-coded according to their lineage of origin, either neutral or dominant. Middle panel: cells are color-coded according to their assigned cluster. Right panel: proportion of afatinib-tolerant and afatinib-sensitive cells within each cluster. (**B-D**) Same as (A) but for day 3, day 6 and day 9 respectively.

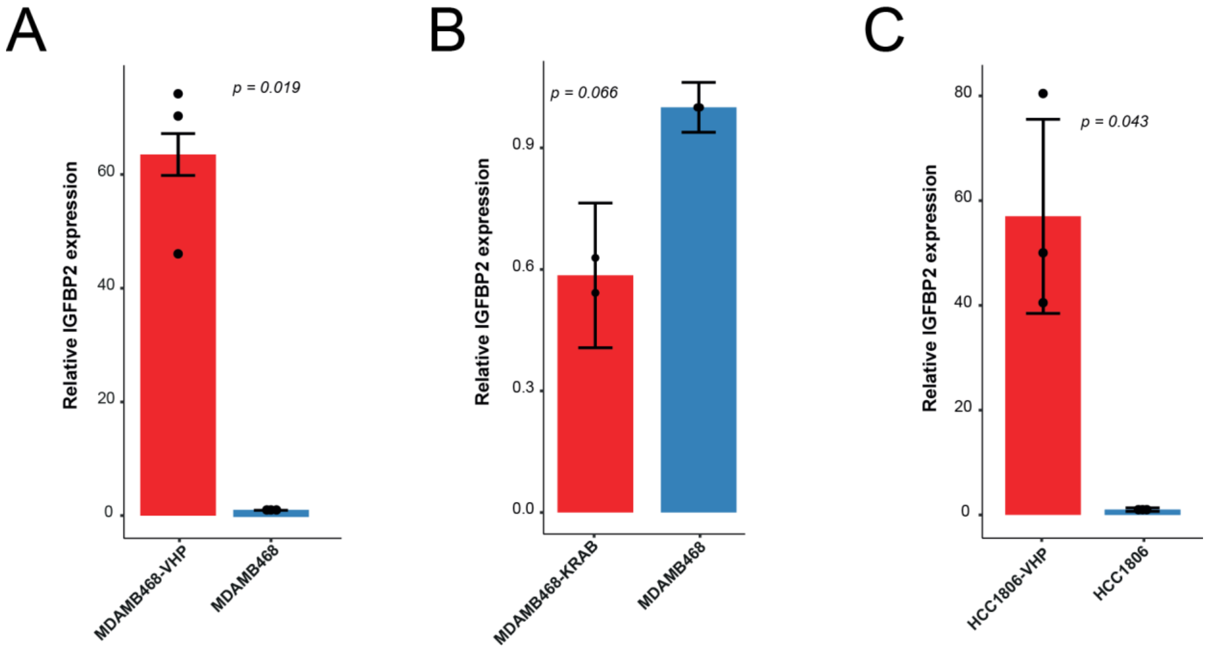


**Supplementary Figure 07 –IGFBP2 expression quantification in CRISP/Cas9 engineered cell lines. (A)** IGFBP2 relative expression in MDA-MB-468 cells with stable overexpression (red) and in control cells (blue). **(B)** Bar plot on IGFBP2 relative expression in both MDA-MB-468 cells with a stable knockdown of the gene (red) and in control cells (blue). **(C)** Relative IGFBP2 expression in both HCC1806 cells stably overexpressing IGFBP2 (red) and in control cells (blue).

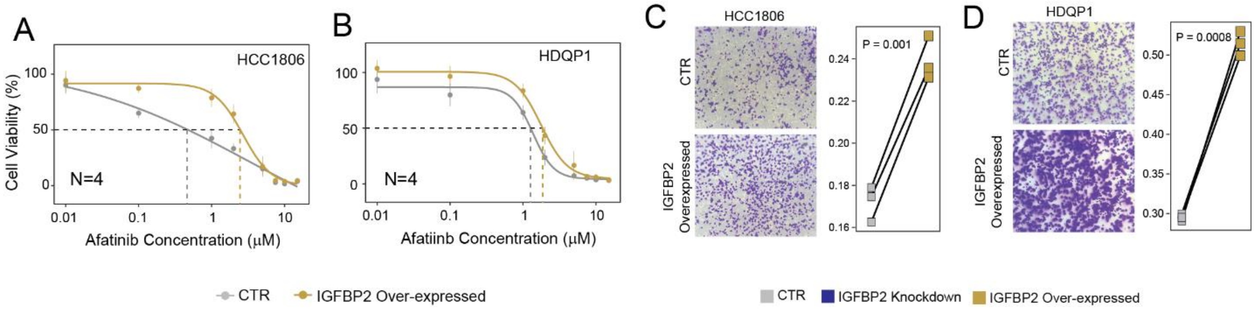


**Supplementary Figure 08 – IGFBP2 implication in Afatinib resistance. (A)** Dose-response curve in terms of cell viability following treatment with Afatinib at the indicated concentrations on HCC1806 IGFBP2 overexpressing cells and relative control cells. **(B)** Dose-response curve in terms of cell viability following treatment with Afatinib at the indicated concentrations on HDQ-P1 IGFBP2 overexpressing cells and relative control cells. **(C)** Transwell migration assay of HCC1806 IGFBP2 overexpressing cells. **(D)** Transwell migration assay of HDQ-P1 IGFBP2 overexpressing cells.

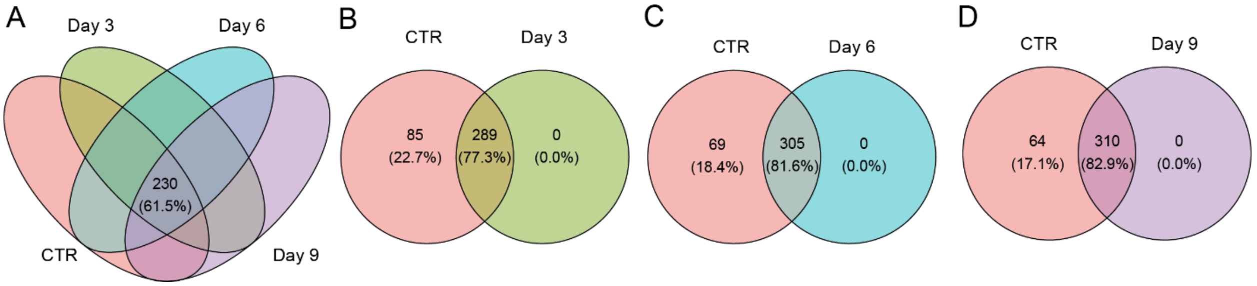


**Supplementary Figure 09 – Differential expression of 374 marker genes of afatinib response across time. (A)** Venn diagram depicting the number of differentially expressed marker genes at each time point compared to untreated Day 0 cells. (**B-D**) Breakdown of shared DEGs of afatinib response across the single timepoints.

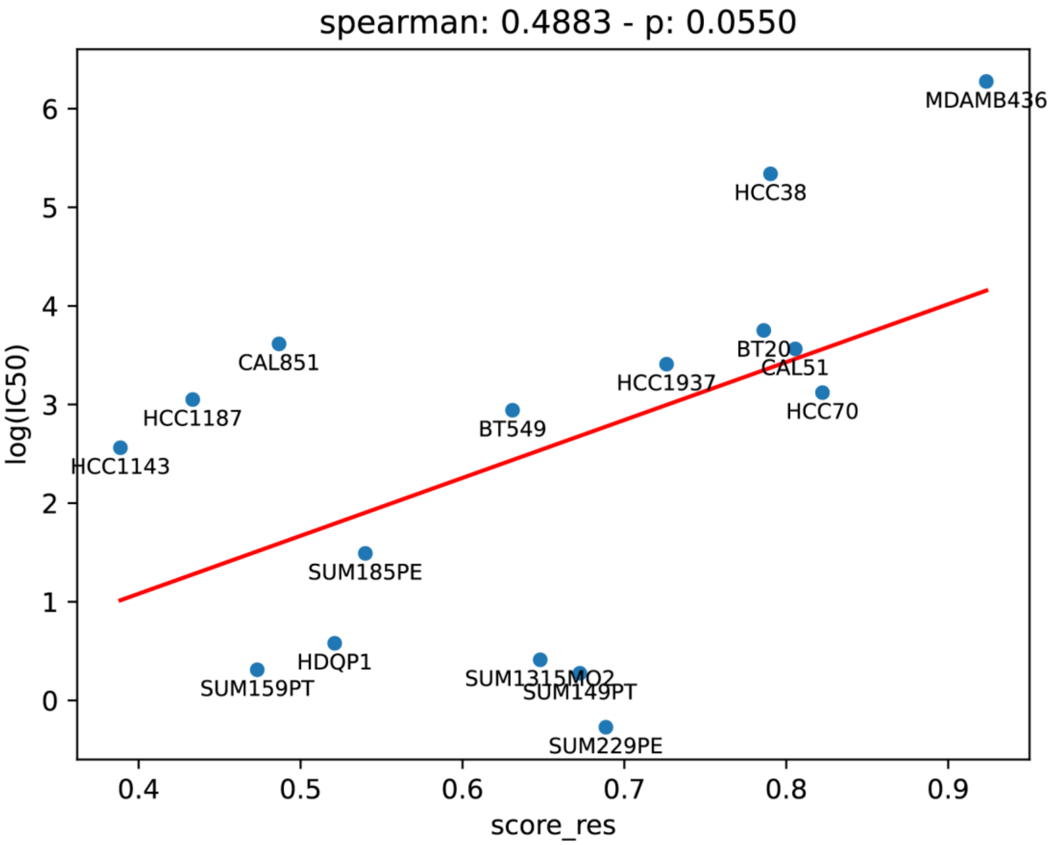


**Supplementary Figure 10 – scASTRAL evaluation of bulk data. (A)** Spearman Correlation Coefficient (SCC) between scASTRAL-predicted afatinib sensitivity and experimentally determined IC50 values for the 16 TNBC cell lines of Figure 5A. The cell lines were first aggregated into pseudobulks (Methods).
